# Functions of ventral visual cortex after bilateral hippocampal loss

**DOI:** 10.1101/673228

**Authors:** Jiye G. Kim, Emma Gregory, Barbara Landau, Michael McCloskey, Nicholas B. Turk-Browne, Sabine Kastner

**Author notes:** **Corresponding author:** Sabine Kastner, Princeton Neuroscience Institute, Washington Road, Princeton University, Princeton, NJ 08544.

## Abstract

Repeated stimuli elicit attenuated responses in visual cortex relative to novel stimuli. This adaptation phenomenon can be considered a form of rapid learning and a signature of perceptual memory. Adaptation occurs not only when a stimulus is repeated immediately, but also when there is a lag in terms of time and other intervening stimuli before the repetition. But how does the visual system keep track of which stimuli are repeated, especially after long delays and many intervening stimuli? We hypothesized that the hippocampus supports long-lag adaptation, given that it learns from single experiences, maintains information over delays, and sends feedback to visual cortex. We tested this hypothesis with fMRI in an amnesic patient, LSJ, who has encephalitic damage to the medial temporal lobe resulting in complete bilateral hippocampal loss. We measured adaptation at varying time lags between repetitions in functionally localized visual areas that were intact in LSJ. We observed that these areas track information over a few minutes even when the hippocampus is unavailable. Indeed, LSJ and controls were identical when attention was directed away from the repeating stimuli: adaptation occurred for lags up to three minutes, but not six minutes. However, when attention was directed toward stimuli, controls now showed an adaptation effect at six minutes but LSJ did not. These findings suggest that visual cortex can support one-shot perceptual memories lasting for several minutes but that the hippocampus is necessary for adaptation in visual cortex after longer delays when stimuli are task-relevant.

## Introduction

Repeated visual stimuli elicit weaker responses in visual cortex than novel stimuli. This adaptation phenomenon (also called repetition suppression or repetition attenuation) can be considered a form of rapid learning and a signature of perceptual memory for previously viewed stimuli. Adaptation has been observed in many visual regions including the lateral occipital cortex (LOC) for repeated presentations of objects (e.g., 1-5) and in the parahippocampal place area (PPA) for repeated presentations of scenes 6-8). Adaptation occurs not only when a stimulus is repeated immediately (with no intervening stimuli), but also after a time lag of several minutes or longer during which intervening stimuli are presented (e.g., 9-14). Although immediate adaptation may reflect a refractory period caused by temporary physiological changes in the local visual area being stimulated (e.g., synaptic depression; see 15), the neural mechanisms underlying long-lag adaptation remain unknown.

For repetitions outside an immediate refractory period, adaptation requires more durable memories for previous stimuli. The memories could be stored in the adapting area itself (i.e., a long-term form of the local changes underlying immediate adaptation), or in other brain regions that provide feedback to the area. The first possibility is called into question by the observation that adaptation can occur for stimuli that have only been seen once before, because cortical learning processes may not support one-shot, long-term memory formation. Indeed, cortex is thought to learn long-term memories only gradually after many exposures and opportunities for consolidation (e.g., 16). Moreover, adaptation can occur after numerous, often highly similar, intervening stimuli, and these stimuli would interfere with sensory representations of the prior stimuli.

For these reasons, here we evaluate the second possibility — that long-lag adaptation in visual cortex is supported by memories stored elsewhere. We focus on one particular memory system — the hippocampus. The hippocampus is a good candidate for supporting adaptation because (a) it is positioned at the top of the ventral visual stream, with anatomical connections from and to many visual areas (17); (b) it can learn rapidly from even a single experience (e.g., 16); (c) it can reinstate memories in visual cortex (e.g., 18-20); (d) it can keep multiple similar stimuli distinct because of sparse coding and pattern separation (e.g., 21); (e) it distinguishes between repeated and novel stimuli of different types (e.g., 14, 22) and (f) it has been linked to cortical adaptation for the repetition of associations (23-24).

We asked whether the hippocampus is necessary for long-lag adaptation, and in particular what role it plays in cortical adaptation for individual stimuli. To establish necessity, we examined patient LSJ (25-28) who has complete bilateral hippocampal loss (Figure 1A). We hypothesized that if the lag between repetitions extends beyond the timescale at which local physiological changes can produce (immediate) adaptation, LSJ will differ from healthy control participants and fail to show adaptation in visual cortex due to the lack of hippocampal feedback. Alternatively, if LSJ shows similar adaptation effects as controls, it is possible that visual cortex can keep track of visual information over certain delays independent of the hippocampus. Because the timescale hypothesized to require the hippocampus is unknown, we varied the repetition lag across several experiments: immediate (in a block design), 30 seconds, 3 minutes, and 6 minutes (all in event-related designs).

**Figure 1.**
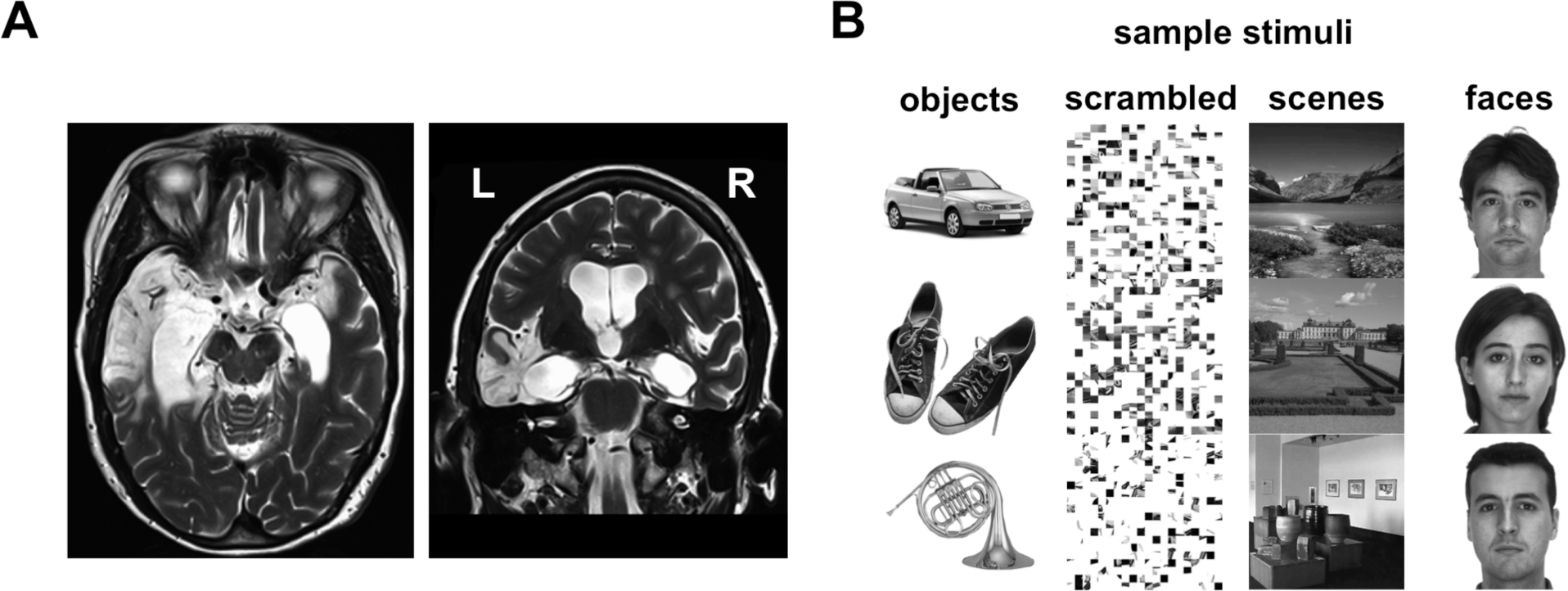
LSJ and localizer stimuli. (A) T2-weighted MRI scan of LSJ’s brain reveals lesions in the bilateral medial temporal lobes (in white), extending laterally to the anterior temporal lobe especially in the left hemisphere. More than 98% of her hippocampus was destroyed bilaterally (27). See Experimental Procedures for further details on the case history. (B) Sample stimuli from the functional localizers used to define object- and scene-selective ROIs. LOC was defined by the contrast of objects vs. scrambled. PPA was defined by the contrast of scenes vs. faces.

Another factor that might affect hippocampal involvement is whether the repeated stimuli are attended and task-relevant. Attention modulates the hippocampus (29), which in turn determines what it learns (30-32). Moreover, attention influences adaptation in visual cortex by increasing selectivity or specificity of the neural population representing the attended sitmuli (33-36). Thus, for the longest lag (6 minutes), which we expected to be most likely to require the hippocampus, we ran two experiments, one with attended and one with non-attended stimuli.

## Results

### Visual Selectivity

To examine adaptation effects in visual cortex, we used functional localizers to define regions of interest (ROIs) for object-selective LOC and scene-selective PPA in each participant. In alternating blocks of trials, LSJ and age-matched controls (total of n = 18 across all experiments) passively viewed series of either objects vs. scrambled objects, or scenes vs. faces (Figure 1B). LOC was defined based on the contrast of greater blood oxygen level-dependent (BOLD) responses to objects than scrambled objects. PPA and other scene-selective areas (transverse occipital sulcus, TOS; retrosplenial cortex, RSC) were defined based on the contrast of greater BOLD responses to scenes than faces. As control regions, retinotopic early visual areas (V1-V4) and the face-selective fusiform face area (FFA) were defined for each participant. Retinotopic areas were defined using a standard topographic mapping procedure from a separate scanning session (8, 37), and FFA was defined based on the contrast faces vs. scenes.

Figure 2 depicts the locations and BOLD time courses of LOC and PPA for LSJ and a representative control participant C1 for visualization purposes. The statistical comparison of LSJ to the control group for this and subsequent analyses was done with a modified independent samples two-tailed *t*-test (38), which accounts for the limited size of the control group and tests the null hypothesis that the single case (LSJ) comes from a population of controls (see Experimental Procedures). Because ROIs were functionally localized in every subject, the comparisons of ROIs between LSJ and controls below are collapsed across experiments. The sizes of LOC and PPA did not differ reliably between LSJ and controls, (*t*_17_ = −1.32, p = 0.22 and *t*_17_ = −1.36, p = 0.20, respectively). The sizes of other visual ROIs also did not differ between LSJ and controls (V1: *t*_17_ = −0.27, p = 0.79; V2: *t*_17_ = 1.31, p = 0.22; V3: *t*_17_ = 0.77, p = 0.50; V4: *t*_17_ = 0.67, p = 0.52; TOS: *t*_17_ = −0.03, p = 0.98; RSC: *t*_17_ = −1.64, p = 0.13; FFA: *t*_17_ = - 0.97, p = 0.35). Despite extensive MTL damage, LSJ showed intact object selectivity in LOC and scene selectivity in PPA and none of the functionally localized ROIs differed in size to that of controls. These results suggest that MTL is not necessary for the basic function and organization of category-selective and retinotopic visual cortex.

**Figure 2.**
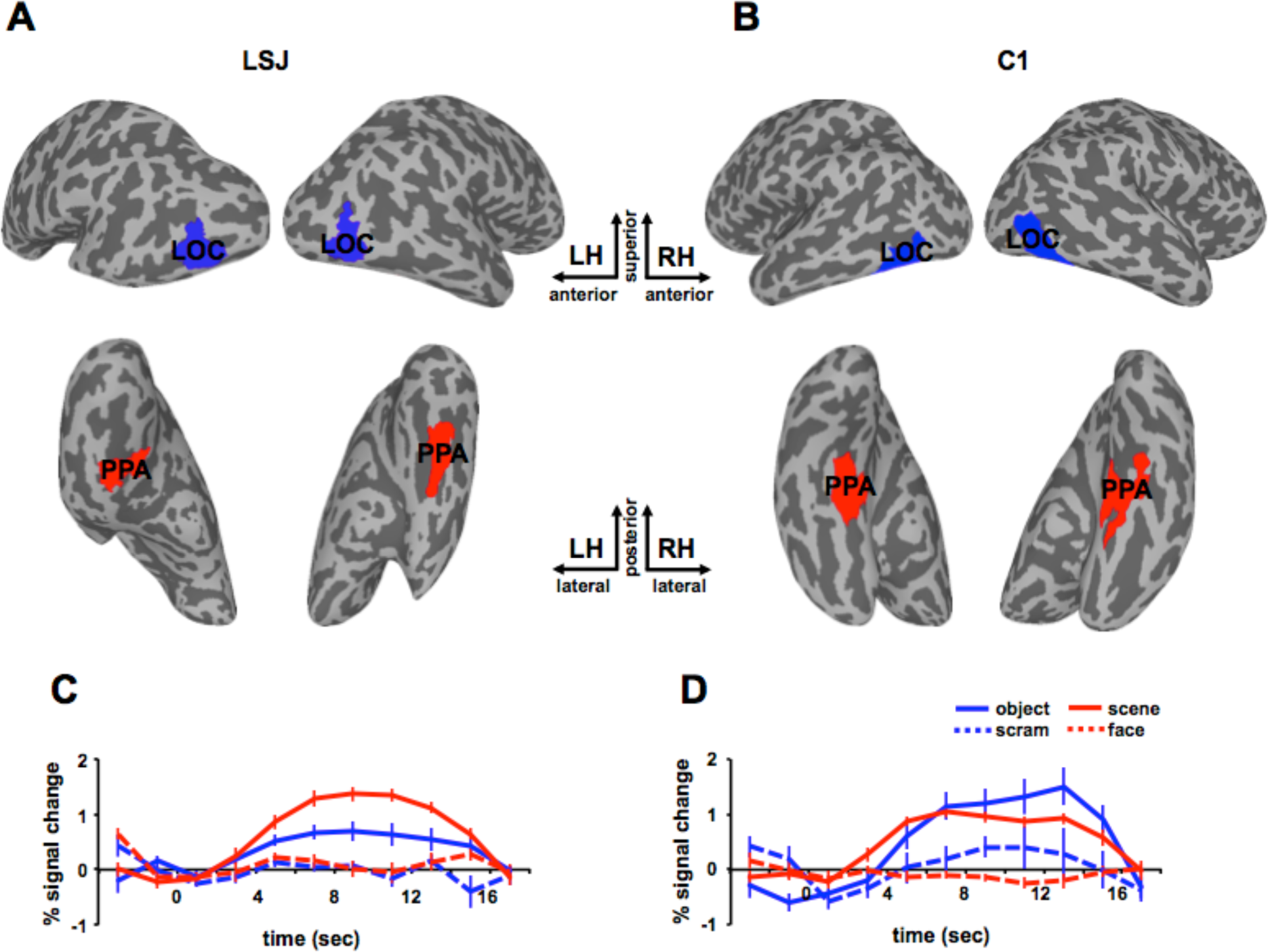
Localizer results. Object-selective LOC (blue) and scene-selective PPA (red) are superimposed on the inflated brains of LSJ (A) and a representative control participant C1 (B). The BOLD time courses in the localizer from (C) LSJ’s and (D) C1’s LOC and PPA are shown for the sake of visualization. The error bars in this figure reflect standard errors across different blocks of trials.

### Immediate Adaptation

In Experiment 1, we used a block design to examine adaptation for immediate stimulus repetitions (Figures 3A and 3B). Previous studies in healthy adults (e.g., 1, 5-6) have shown robust adaptation in LOC and PPA when comparing blocks that contain, respectively, one object or scene presented repeatedly (“repeat”) with blocks of the same total number of stimuli but with each stimulus being novel (“new”). We hypothesized that such short-lag adaptation is mediated by the visual system and thus does not depend on the hippocampus. Consequently, we expected that both LSJ and controls would show adaptation effects in this paradigm. This first experiment was an important step to establish that the adaptation paradigm was a feasible way to test LSJ in an fMRI study, given the difficulties of working with severely amnesic patients in such an environment.

**Figure 3.**
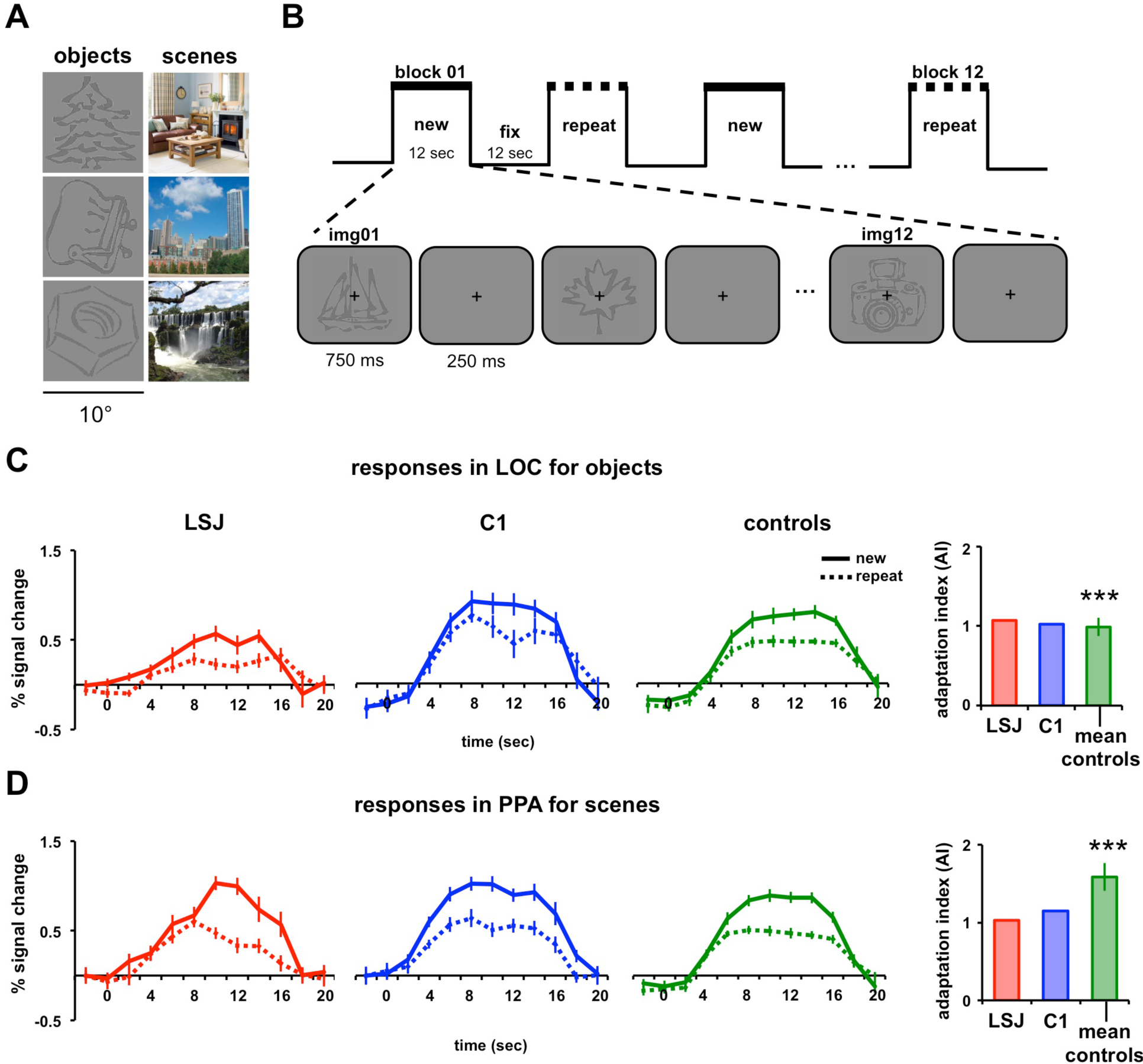
Block design and immediate adaptation. (A) Example line-drawing objects and scenes used as stimuli. (B) Blocks alternated between “new” (12 different stimuli) and “repeat” (1 stimulus repeated 12 times) conditions, with order counterbalanced across runs. Objects and scenes were shown in separate runs. BOLD time courses and AIs for LSJ (red), a representative control C1 (blue), and the average of all controls (green) for (C) objects in LOC and (D) scenes in PPA. The asterisks above the control average denote significant adaptation effects from zero using a one-sample t-test (*** *p* < 0.001). Error bars in the line graphs for LSJ and C1 denote standard errors across blocks of trials. The error bars for average controls (for this and subsequent figures) denote standard errors across subjects. Using the Crawford and Howell’s t-test for case-control comparisons (38), LSJ’s AIs did not reliably differ from control AIs for both LOC and PPA.

Figures 3C and 3D show the BOLD time courses in LOC for objects and PPA for scenes from LSJ, C1, and averaged controls (n = 8). As expected, control participants showed greater BOLD responses to the new vs. repeat blocks. To quantify these effects, we computed, for each ROI in each participant, an adaptation index (AI): the difference in peak responses to the new and repeat conditions as a proportion of the variance across blocks from the combined conditions (see Experimental Procedures). AI values greater than zero denote greater BOLD responses for the new than repeat conditions. Controls had AIs reliably greater than zero for objects in LOC (*t*_7_ = 8.58, p < 0.0001) and scenes in PPA (*t*_7_ = 8.90, p < 0.0001). LSJ also showed positive AIs for both objects in LOC and scenes in PPA. Furthermore, LSJ’s AI for LOC (*t*_7_ = 0.21, p = 0.84) and PPA (*t*_7_ = −1.04, p = 0.33) did not differ reliably from controls (see Figure S1 to compare LSJ’s AI values against individual controls for this and subsequent Exps.). See Figure S2 and Tables S1-S2 for adaptation effects in other ROIs (for this and subsequent Exps.). To examine whether or not LSJ’s adaptation effects are expected by chance, we performed a permutation test to compute the null distribution of AIs by randomizing the condition labels and computing AIs across 1000 iterations (see Figure S3). This analysis showed that LSJ’s AIs for LOC and PPA (for this and subsequent Exps. where LSJ showed adaptation effects) was not due to chance and well below the 5% tail of the permutation distribution.

Neither controls nor LSJ showed hemispheric differences in AIs for this and subsequent experiments (see Figure S4 and Tables S3-S4). Consistent with our hypothesis, these results suggest that the hippocampus is not necessary for immediate adaptation effects in ventral visual cortex. It also demonstrated the feasibility to test the patient in this type of study.

### Long-Lag Adaptation Without Attention

In Experiment 1, at the boundary condition of immediate repetitions (i.e., zero lag), it appeared that local physiological changes within visual cortex were sufficient for adaptation, or at least that such adaptation could occur independently of the hippocampus. However, these physiological changes may dissipate some period of time after stimulus presentation, which might correspond to the lag at which longer-term memories are required and adaptation would become hippocampally dependent. As we did not know *a priori* at which timescale this might happen, we manipulated lag parametrically across the remaining experiments.

#### 30-s Lag

Experiment 2 used a rapid event-related design in which each stimulus was repeated once at a lag of approximately 30 s or 6 intervening stimuli (Figure 4A). We again examined BOLD responses to objects in LOC and scenes in PPA (in separate runs). While passively viewing the stimuli, participants engaged in an “alphabet game” at the center of the screen that required them to think of words that began with a cued letter (see Experimental Procedures). This task was chosen because it is one of LSJ’s favorite activities and it reduced attention to the object and scene stimuli, as they were task-irrelevant and presented in the background of the letters.

**Figure 4.**
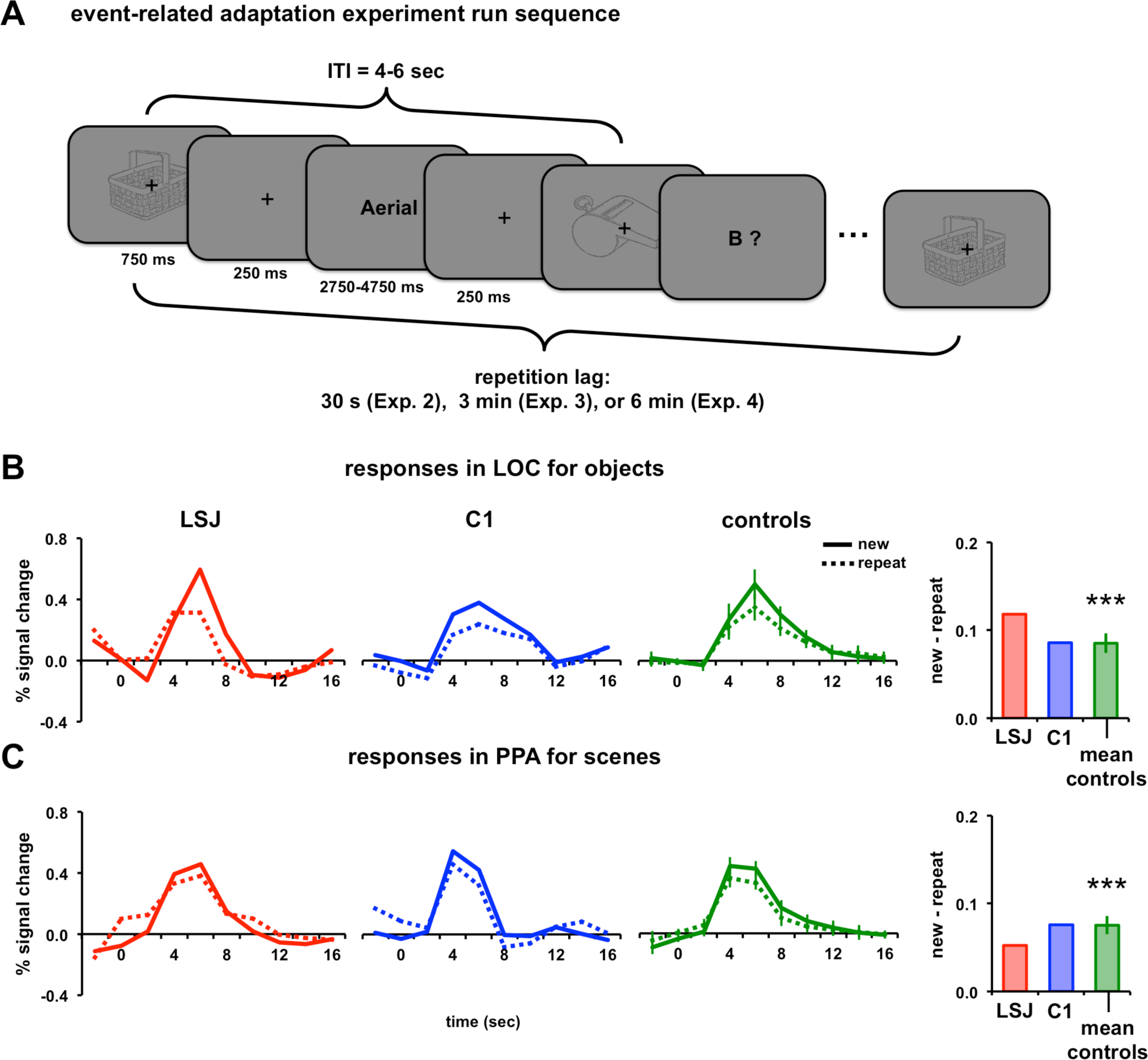
Event-related design and 30-s lag adaptation. (A) Example sequence for the lagged adaptation experiments with rapid-event related design. Each stimulus was repeated just once, after 30 s (Experiment 2), 3 mins (Experiment 3), or 6 mins (Experiment 4) on average. Participants performed a demanding alphabet task interleaved with the stimuli, in which the computer generated a word starting with one letter of the alphabet and then the participant generated a word covertly starting with the next letter. BOLD timecourses and AIs for repetitions after a 30-s lag of (B) objects in LOC and (C) scenes in PPA. For calculating AIs, peak responses were defined as the average BOLD response 4-8 s post-stimulus.

Figures 4B and 4C show the BOLD time courses and AIs for LSJ, C1, and the control group, averaged for object presentations in LOC and scene presentations in PPA. The AI quantification for this experiment (and all subsequent event-related experiments) was the difference between the average peak responses (time points 4-8 s post stimulus onset) for the new and repeat conditions (see Experimental Procedures). The controls’ AIs for objects in LOC were reliably positive (*t*_7_ = 7.76, p < 0.001), indicating adaptation approximately 30 s after first exposure to a stimulus. LSJ also showed a positive AI, and it did not differ from controls (*t*_7_ = 1.00, p = 0.35). Likewise, the controls’ AIs for scenes in PPA were reliably positive (*t*_7_ = 7.34, p < 0.001). LSJ also showed a positive AI, which again did not differ from controls (*t*_7_ = - 0.74, p = 0.48).

Similar to immediate adaptation, both controls and LSJ showed reliable adaptation in LOC and PPA, suggesting that the hippocampus is not necessary to bridge across repetitions spaced by 30 s and that cortex keeps track of visual input over this period of time. It is notable that these effects occurred despite the fact that attention was drawn away from the stimuli to a different task, suggesting a highly automated tracking of stimuli by the visual system across this time interval.

#### 3-min Lag

Having observed reliable adaptation in both LSJ and controls at a 30-s lag, in Experiment 3, we increased the lag to approximately 3 mins or 42 intervening stimuli. Anecdotally, LSJ seems to have a shorter time window of memory than this (e.g., she repeats questions, and forgets having done tasks, after 1-2 mins), and so this lag seemed like a reasonable timescale for possibly observing engagement of the hippocampus. Everything else was the same as Experiment 2, except that the longer lag meant that the initial and repeated presentations of each stimulus occurred in different runs. LSJ and controls passively viewed the stimuli shown in the background while engaging in the alphabet game at the center of the screen.

Figures 5A and 5B show the BOLD time courses and AIs for LSJ, C1, and the control average for objects in LOC and scenes in PPA. The control AIs for objects in LOC were again reliably positive (*t*_7_ = 5.04, p < 0.002) and not different from LSJ’s AI (*t*_7_ = −0.12, p = 0.91). Likewise, the control AIs for scenes in PPA were again reliably positive (*t*_7_ = 4.89, p < 0.002) and not different from LSJ’s AI (*t*_7_ = 1.51, p = 0.18). Thus, the pattern of results for a 3-min lag was very similar to that for the 30-s lag.

**Figure 5.**
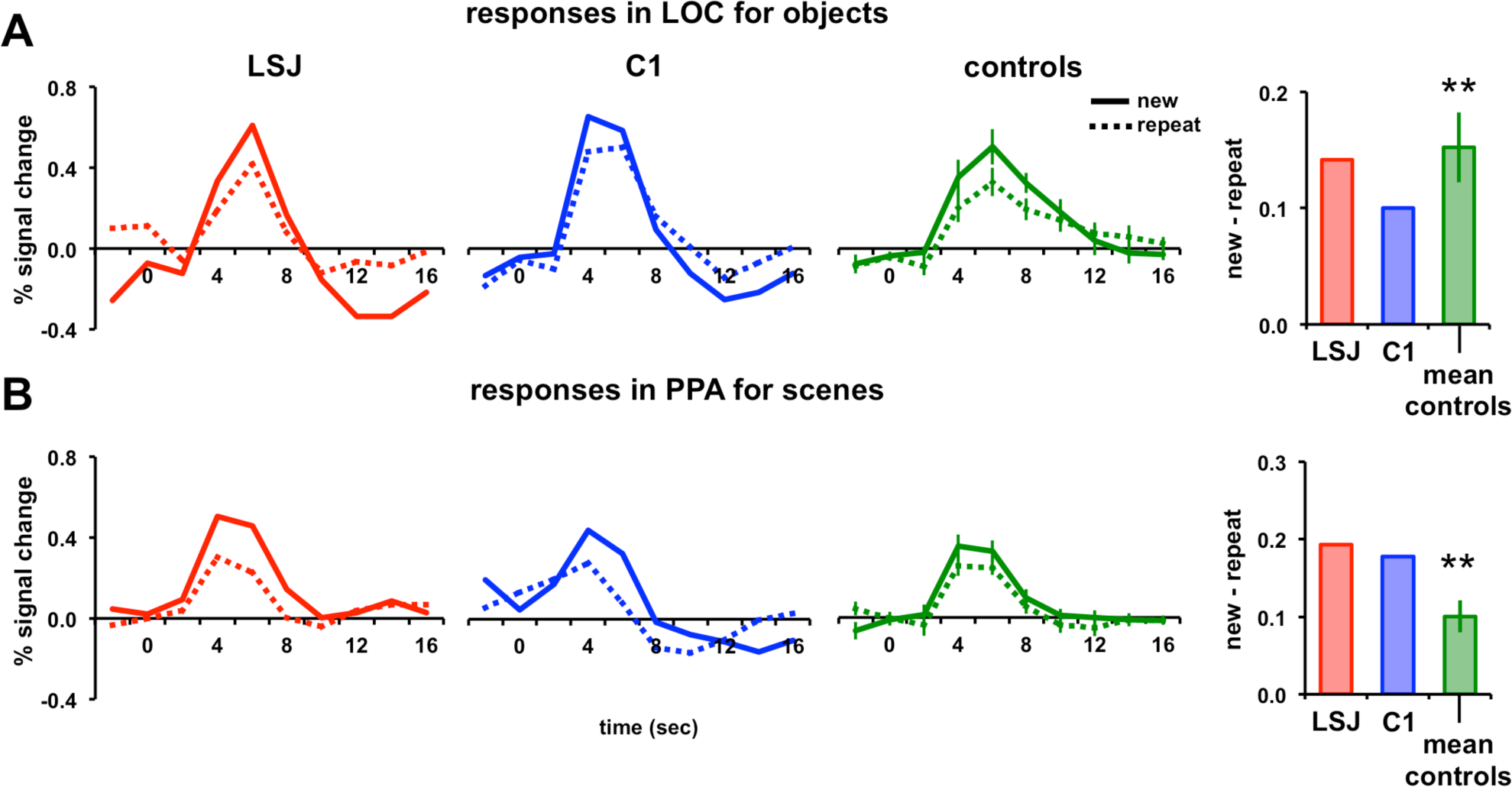
3-min lag adaptation. BOLD timecourses and AIs for repetitions after a 3-min lag of (A) objects in LOC and (B) scenes in PPA (** p < 0.01).

#### 6-min Lag

Given that increasing the lag from 30 s to 3 mins did not impact the results, in Experiment 4 we doubled it again to approximately 6 mins or 68 intervening stimuli.

Figures 6A and 6B show the BOLD timecourses and AIs for LSJ, C1, and the control average for objects in LOC and scenes in PPA, respectively. In contrast to the prior experiments, the increased lag eliminated the long-lag adaptation effect observed previously. The control AIs did not reliably differ from 0 for either objects in LOC (*t*_7_ = 0.67, p = 0.52) or scenes in PPA (*t*_7_ = 1.06, p = 0.33). Using an independent samples t-test, the AIs for the 6-min lag were reliably weaker than for the 30-s lag in both LOC and PPA (*t*_14_ = 3.66, p = 0.003 and *t*_14_ = 3.91, p = 0.002, respectively) and compared to the 3-min lag (*t*_14_ = 4.06, p = 0.001 and *t*_14_ = 3.66, p = 0.003, respectively). LSJ was not different from controls for either objects in LOC (*t*_7_ = −0.41, p = 0.69) or scenes in PPA (*t*_7_ = −0.88, p = 0.41).

**Figure 6.**
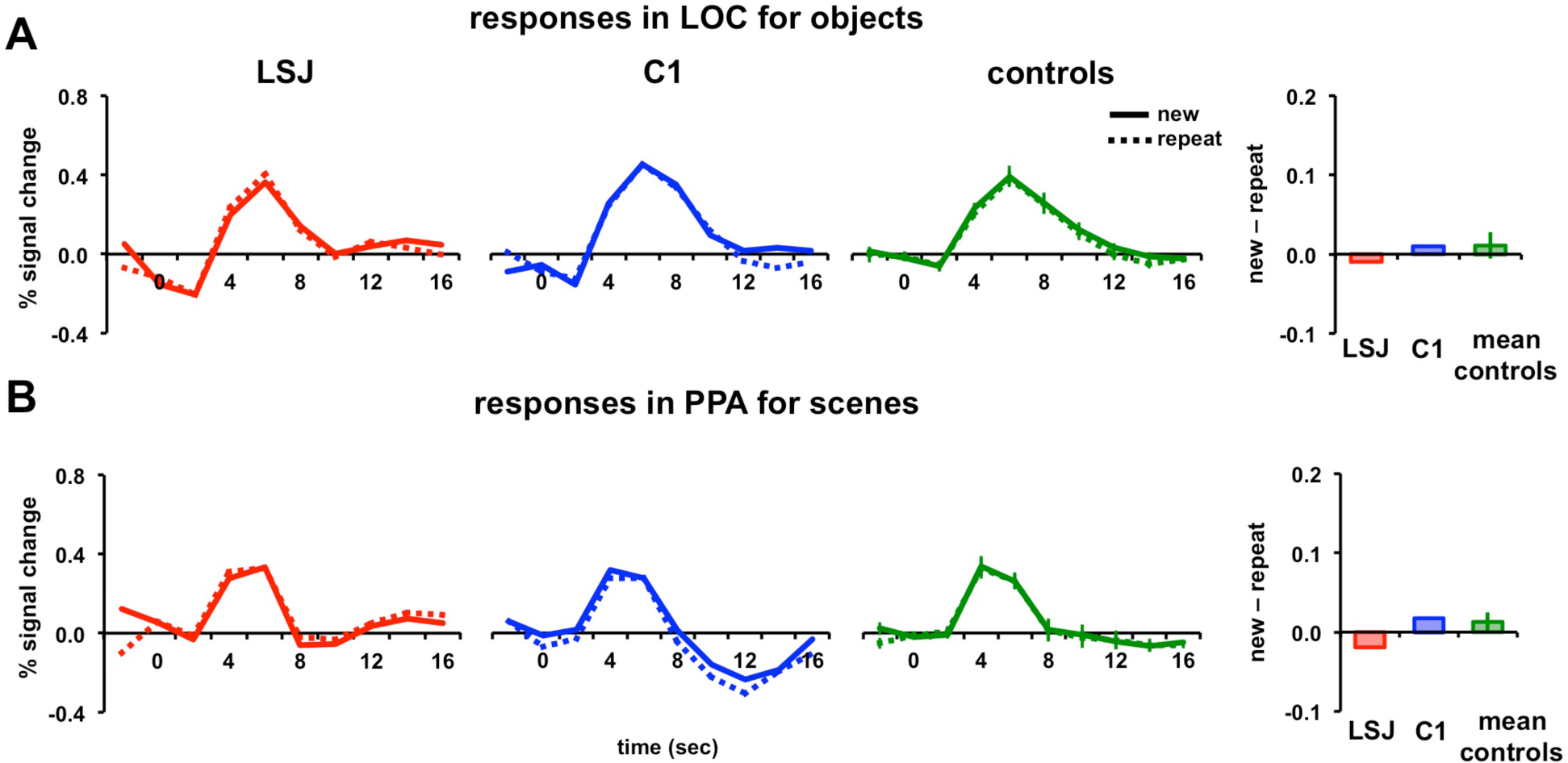
6-min lag adaptation. BOLD timecourses and AIs for repetitions after a 6-min lag of (A) objects in LOC and (B) scenes in PPA.

Although LSJ no longer showed adaptation, the lack of adaptation in controls prevents us from concluding based on this experiment that the hippocampus is necessary. Instead, it appears that the automatic tracking of stimulus information performed by the visual system occurs on the order of a few minutes, but reaches a limit within a 3-6 minute time frame.

### Long-Lag Adaptation With Attention

Why did we not observe adaptation in controls after a 6-min lag, despite the fact that previous studies (9-10, 39) have found such effects at even longer lags? One difference is that the prior studies employed behavioral tasks that required participants to attend to and make categorization judgments about the stimuli. For instance, participants might be instructed to name objects (e.g., 39, 9-10) or perform a categorization task such as deciding whether an object is bigger or smaller than a shoebox (12, 40-41). In contrast, Experiments 2 - 4 used a task that was orthogonal to the repeating stimuli and thus drew attention away from them. Consistent with the importance of stimuli being task-relevant, selective attention has been shown to modulate long-lag adaptation (34, 36, 42-43).

To examine the role of attention in long-lag adaptation as well as possible interactions with hippocampus, in Experiment 5 we asked participants to perform a go/no-go categorization on the stimuli (Figure 7A). All other experimental parameters were identical to Experiment 4 (e.g., lag was 6 minutes). For objects, participants were instructed to press a button if the presented object was a natural object; and for scenes, if the presented scene was an indoor scene. To help LSJ remember the task, a task prompt (“Press a button if this is a natural object” for objects and “Press a button if this is an indoor scene” for scenes) was shown on top of the screen throughout the experimental runs for all subjects.

**Figure 7.**
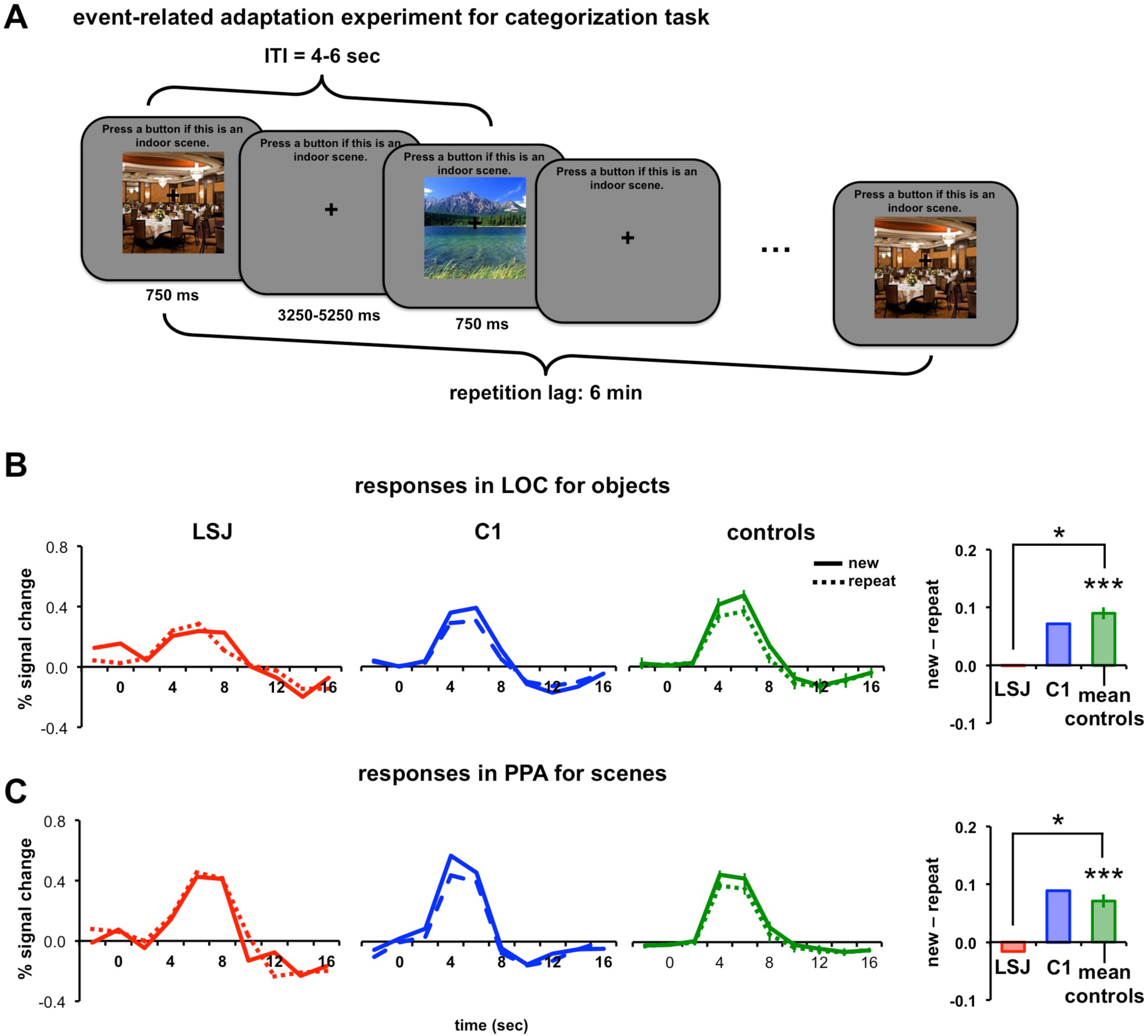
Task-relevant design and 6-min lag adaptation. (A) Example trial sequence using an event-related design with a categorization task. For scenes (shown here), participants were instructed to press a button if the presented scene was an indoor scene. For objects, participants were instructed to press a button if the presented object was a natural object. The task prompt was shown on the top of the screen throughout the experiment to minimize the memory demands of the task on LSJ. BOLD time courses and AIs for LSJ, C1, and the average of all controls for (B) objects in LOC and (C) scenes in PPA. (* p < 0.05).

In contrast to Exp. 4 controls showed reliable adaptation effects (Figure 7B and 7C) in LOC for objects (*t*_7_ = 8.38, p < 0.0001) and PPA for scenes (*t*_7_ = 6.26, p < 0.001). Strikingly, LSJ did not show any adaptation, with AIs near 0 and lower than all controls for objects in LOC (*t*_7_ = −2.83, p < 0.03) and scenes in PPA (*t*_7_ = −2.56, p < 0.04). See Figure S1 for comparison of LSJ and individual controls’ AI values. These findings suggest that the hippocampus is necessary for tracking stimuli over a time period of 6 minutes after initial presentation when the repeating stimuli are task-relevant. The comparison of AIs across experiments for controls and LSJ are summarized in Figure 8.

**Figure 8.**
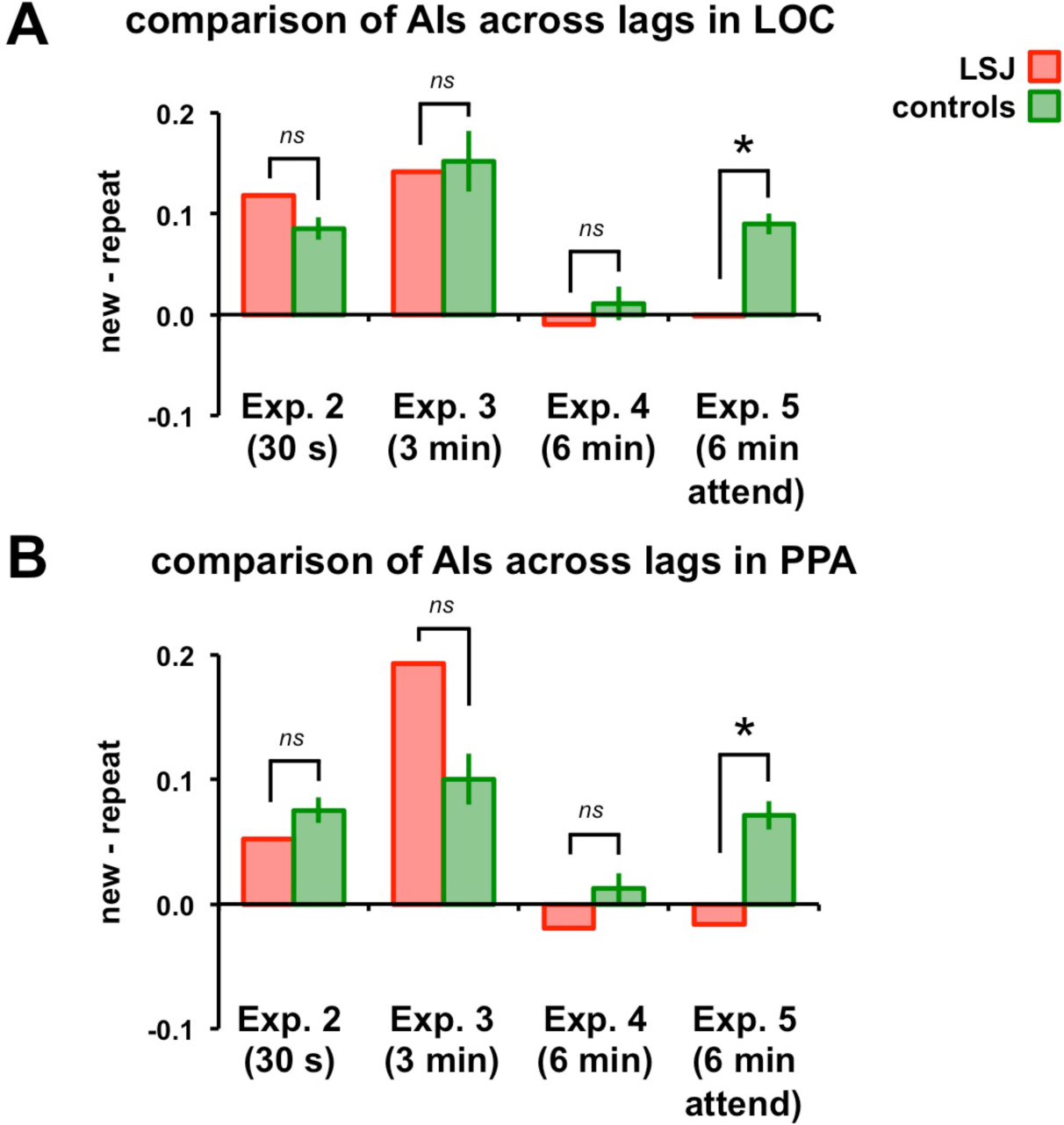
Comparison across experiments. AIs from Experiments 2-5 are re-plotted for LSJ (in red) and controls (in green) for objects in LOC (A) and scenes in PPA (B).

Behavioral accuracy in the tasks was high for both LSJ (88.6% for objects and 89.7% for scenes) and controls (mean = 94.6% for objects and 94.2% for scenes), and LSJ’s accuracies did not differ reliably from those of controls (*t*_7_ = −1.19, p = 0.27 for objects and *t*_7_ = −1.06, p = 0.33 for scenes). Response times (RTs) did differ between LSJ and controls for objects (789 vs. mean of 593 ms, respectively; *t*_7_ = 2.96, p = 0.02), but not for scenes (687vs. mean of 610 ms, respectively; *t*_7_ = 1.20, p = 0.27).

Making the stimuli task relevant also let us examine response priming in behavior by comparing RTs for the first vs. second exposure of each stimulus. Controls were reliably faster for the second than the first exposure of objects (mean difference = 43.8 ms, *t*_7_ = 3.90, p < 0.006) and marginally faster for scenes (mean difference = 10.9 ms, *t*_7_= 2.03, p < 0.08). LSJ also showed a numerical benefit for second vs. first exposures (8.1 ms for objects and 10.9 ms for scenes), and these differences were not reliably different from controls (*t*_7_ = −1.06, p = 0.32 and *t*_7_ = −0.32, p = .76, respectively). Controls did not exhibit greater accuracy on the categorization tasks for the second vs. first exposure of objects (mean difference = −0.44%, *t*_7_ = 0.91, p = .39) or scenes (mean difference = 0.21%, *t*_7_ = −0.60, p = 0.56). LSJ showed similar accuracy results (difference of 0.41% for objects, *t*_7_ = 0.58, p = 0.57 and 0.22% for scenes, *t*_7_ < 0.01, p > 0.99).

Together, these results suggest that LSJ was able to perform the behavioral tasks. The only reliable difference from controls was the overall RT in the object task, and thus any adaptation effects that hold for both objects and scenes cannot be fully explained by differences in task performance or difficulty. The behavioral priming for controls and LSJ is consistent with prior studies showing preserved long-lasting behavioral priming in hippocampal amnesia (44-46). Moreover, the presence of this behavioral priming effect does not entail that adaptation occurred in visual cortex, as behavioral response priming has been shown to be supported by frontal and other structures also preserved in LSJ (41, 47-48).

## Discussion

We conducted a case study of LSJ, a patient with complete bilateral hippocampal loss, across a series of five fMRI experiments with two sessions per experiment (to replicate the findings across object and scene stimulus classes). Our findings provide a comprehensive account of the role of the hippocampus in neural adaptation (summarized in Figure 8). Using localizers, we first defined ROIs for object-selective LOC and scene-selective PPA, as well as other category-selective (TOS, RSC, FFA) and early visual areas (V1-V4). None of these ROIs differed in size from controls, suggesting that the organization and selectivity of ventral visual cortex does not depend on the hippocampus. Also like controls, LSJ showed adaptation in LOC and PPA for object and scene repetitions, respectively, whether they occurred immediately, after 30 s, or after 3 mins. LSJ did not show adaptation for repetitions after 6 mins, regardless of whether the stimuli were task-relevant or -irrelevant. In contrast, controls showed robust adaptation after 6 mins when the stimuli were task-relevant. These results suggest that visual cortex, independent of a contribution from the hippocampus, is able to sustain perceptual memory for stimuli viewed once up to about 3 minutes but not 6 minutes ago. Attention can modulate adaptation at long (at least 6 minutes) lags, and the hippocampus plays a vital role in mediating long-lag adaptation for task-relevant stimuli in visual cortex. Although we do not currently have a detailed mechanistic account of our results, below we provide interpretations that can pave the way for future studies.

It has previously been argued that distinct neural mechanisms underlie immediate and long-lag adaptation effects (e.g., 9,10,12,49). It is possible that immediate adaptation effects are driven by local mechanisms within visual cortex that do not require feedback from higher-order areas, but instead reflect physiological changes such as synaptic depression or neuronal fatigue (e.g., 15). EEG studies (12, 49) have supported such a distinction by demonstrating different temporal components of adaptation: an early signature (150-300 ms after stimulus onset) and a late signature (400-600 ms after stimulus onset). Accordingly, the early signature may reflect local physiological changes, whereas the late signature may reflect feedback from higher-order areas that maintain information about recent stimuli. Our results can be interpreted in this framework by positing that repetitions up to three minutes apart produce adaptation in the early component, but that beyond this lag, adaptation would appear in a late component. Moreover, LSJ and other hippocampal patients should show adaptation only in the early component. Future studies examining the temporal dynamics of adaptation effects would be needed to test this hypothesis with our stimuli, tasks, and lags.

Putting aside the longest lag, it appears surprising that adaptation could occur in visual cortex after three minutes without a hippocampus. Indeed, adaptation in LSJ at this lag was robust for objects in LOC and scenes in PPA. Prior fMRI (9, 10, 39) and monkey electrophysiology studies (50-52) have shown adaptation effects at long lags. However, to what extent these effects were dependent on the hippocampus or other structures in the medial temporal lobe was unclear. Here, we show that without the hippocampus, LOC and PPA can track visual information over the course of three minutes. Recently, it has been suggested that LOC and PPA have temporal receptive windows of several minutes (53). The temporal receptive window is mapped with a stimulus that has a higher order structure (i.e., a movie stimulus with a continuous plot) and temporal receptive window would be a representation of some aspect of a stimulus that bridges over a long time. The stimuli in our experiments were random with no overarching structure of the objects or the order in which the objects were presented. Thus, it is unlikely that the temporal receptive window can explain the adaptation effects observed in LOC and PPA for longer lags up to 3 minutes. One alternative is that working memory could play a role in supporting adaptation effects up to 3 minutes. This also seems unlikely because our stimuli consisted of hundreds of images that were presented in the background when subjects were performing an orthogonal task. The alphabet-naming task diverted their attention away from the stimuli, discouraging their active maintenance in working memory during the experiment.

Even more strikingly, LSJ seemed to show adaptation at 3 minutes lag in early visual areas (V1-V4), especially for scenes (Figure S1 and Table S2). However, whether stimulus information is maintained this long in such areas is unclear, especially because they are typically believed to have a relatively short temporal window over which sensory information is integrated (53). One possibility is that higher-level areas like LOC and PPA track the information and send feedback to early visual areas. Another possibility is that a memory system other than the hippocampus is involved. For example, the intact frontal cortex of LSJ may have automatically maintained recent stimuli up to this lag.

Previous studies of long-lag adaptation have found that the magnitude of adaptation decreases with increasing lag between the first and second presentations of stimuli (11, 14, 54-55). This is consistent with our finding that robust adaptation for task-irrelevant stimuli at the three-minute lag was eliminated by 6 minutes. However, we found that attention was another important factor, as adaptation re-emerged at 6 minutes when the stimuli were made task-relevant. Thus, while adaptation dissipates over time, attention may act to extend it.

We occasionally, although not consistently, observed adaptation effects in regions that are not selective for objects or scenes (e.g., adaptation in PPA or early visual areas to repetition of objects; Figure S2 and Tables S1 and S2). This was more prominent in the shorter lag experiments. These findings are consistent with previous studies showing that adaptation effects can be found in regions without visual selectivity (e.g., 13). Moreover, attention increases selectivity in ventral visual cortex (33), and the adaptation effects observed in controls when stimuli were task-relevant were more specific to the corresponding category-selective ROIs (Figures S2 and Tables S1 and S2).

Could the lack of adaptation in LSJ with a six-min lag when the stimuli were task-relevant (Experiment 5) be explained by differences in task performance or difficulty? Although it is challenging to eliminate this possibility, we think that such factors are unlikely to explain our results in a parsimonious way. First, accuracy did not differ between LSJ and controls in either the object or scene tasks. Second, even though LSJ was slower than controls in the object task, she was not reliably slower in the scene task, and both stimulus categories produced the same pattern of differential adaptation effects. Finally, LSJ showed evoked BOLD responses in LOC and PPA for new stimuli that were comparable to controls, suggesting that she was perceptually processing the stimuli. Together this evidence suggests that LSJ and controls’ performance to the categorization task was qualitatively comparable and whatever differences in adaptation effects between LSJ and controls cannot be attributed to differences in performance difficulties, attentional deployment or deficits.

A central result of our study is that the hippocampus appears to support long-lag adaptation beyond three minutes when stimuli are task-relevant and thus attention and memory systems are interacting. How could the hippocampus support such adaptation? One potential mechanism is that the hippocampus (and potentially other areas) forms long-term memories of the attended stimuli, and such memories are the only remaining trace of the stimuli after six minutes. When a repeated stimulus is subsequently perceived, the memory of the initial presentation is retrieved and reinstated in visual cortex (e.g., 20, 56-57). This reinstatement could have the effect of facilitating sensory processing in the visual cortex, leading to an overall reduction in visual activity. This cannot occur in LSJ, as she lacks these hippocampal memories, potentially explaining why she did not show adaptation at the longest lag. This proposed relationship between hippocampally dependent long-term memory and long-lag adaptation is consistent with prior studies that have linked adaptation to behavioral measures of long-term memory (e.g., 42, 58).

To examine whether or not the hippocampus is involved at all in adaptation, we examined the hippocampus of controls in all of our experiments (see Figure S6). As the repetition lag increased, hippocampus activity changed from no visually evoked response in the immediate to short-lag experiments to evoked responses at 6 min lag experiments for both the object and scene tasks. This pattern of hippocampus activity suggests that the hippocampus is more engaged in the longer than shorter lag adaptation.

Support of the view that the hippocampus is involved in adaptation effects with enhanced attention to stimuli come from studies examining repetition effects on eye movements with amnesic patients with hippocampus damage (59-60). While these patients demonstrate the behavioral repetition effect of eye movement on repeated scenes under passive viewing (i.e., fewer fixations and fewer regions of sampling when viewing repeated versus new stimuli), they do not show this effect when instructed to be aware of the old and new status of the scenes. Our results support these findings that repetition effects with enhanced attentional processing is hippocampus-dependent.

Attention and memory are intricately connected systems and selective attention has been shown to facilitate memory formation (61). The precise neural mechanism whereby attention influences memory is still debated, with recent studies suggesting that attention can modulate hippocampal activity during stimulus encoding (30-31). Selective attention has also been shown to stabilize neural representations in the hippocampus during encoding (32). Thus, the dependence of long-lag adaptation on attention in controls is consistent with a mechanism based on long-term hippocampal memory. Specifically, task-relevant stimuli may have produced more robust memory traces in the hippocampus, which in turn facilitated processing in visual cortex via reinstatement during repetitions. This idea is also consistent with findings that adaptation effects in category-selective cortex interact with episodic memory processes in the hippocampus (24).

Another mechanism for the hippocampal dependence of long-lag adaptation could be based on the role of the hippocampus in predictive coding. Several studies have shown that expected stimuli evoke weaker responses in visual cortex, and this has been provided as an explanation for adaptation (62). Namely, because the environment is relatively stable over time, we are more likely to encounter a recently seen stimulus again than a new stimulus for the first time. Accordingly, repeated stimuli should be more expected than novel stimuli, all else being equal. In predictive coding models, this has the consequence of potentiating or “explaining away” sensory representations of the expected stimuli, preventing these stimuli from producing a net increase in activity. As a result, the visual system preferentially represents unexpected stimuli (i.e., prediction errors), in the service of learning to generate better predictions over time. Where such expectations are stored is the subject of much investigation, with candidates including orbitofrontal cortex (63) and the ventral striatum (64). However, the hippocampus has recently emerged as a candidate system for learning about the structure of the visual environment (65) and for generating predictions about upcoming visual stimuli based on this structure (66). Without a hippocampus, LSJ may not be able to form such expectations as well (27), attenuating their impact on the visual system.

In Experiment 5, both LSJ and controls exhibited behavioral priming in the categorization tasks. This is consistent with previous studies demonstrating preserved behavioral priming in hippocampal patients (44-46). These amnesic patients, despite being severely impaired in recognizing previously presented stimuli, still exhibited long-lasting priming effects to similar degrees as healthy controls (46). Behavioral priming is often accompanied by repetition suppression in ventral visual cortex (e.g., 12, 41, 47-48, 67), but priming can be dissociated from such suppression (68) and is thought to depend instead on frontal cortex (41, 47-48, 67). Indeed, LSJ exhibited behavioral priming effects without adaptation in LOC or PPA.

In summary, the present study offers insights into the role of the hippocampus in the function of ventral visual cortex and the tracking of visual information in cortex without the support of the hippocampus. Although an intact hippocampus does not appear to be required for adaptation from seconds to a few minutes, it may be required for longer delays under conditions of enhanced attentional processing. This further highlights how the hippocampus can contribute to visual perception and learning, justifying and explaining its position at the top of the visual processing hierarchy.

## Methods

### Case History

LSJ is a 68 year-old (62 at the time of the first and 63 at the time of last scan session), right-handed, college-educated woman. She is a highly successful artist and amateur musician. She contracted herpes encephalitis in 2007. High-resolution anatomical MRI revealed that more than 98% of her hippocampus was destroyed bilaterally (27); she also has extensive damage to other MTL and anterior temporal regions, especially in the left hemisphere (Figure 1A). Her medical history prior to this event was unremarkable. LSJ suffers from anterograde and retrograde amnesia and her score on the Wechsler Memory Scale’s General Memory is < 0.1%. Her basic sensory and language abilities seem intact. A thorough examination of LSJ’s memory functions is detailed in Gregory et al.’s study (25; For additional reports concerning LSJ’s memory and learning abilities, see 26-28).

### Participants

A total of 18 age-matched control subjects participated in the experiments (all right-handed, two males, mean age = 62.8 (range 56 - 69), no history of neurological disorder). For each experiment, there were 8 control participants. The same 8 controls participated in Experiments 1-3 and three of these controls also participated in Experiments 4 and 5. The other 10 participants tested in Experiments 4 and 5 were distinct individuals. Because of the extensive nature of the experiments, we were unfortunately unable to recruit the same control participants for all experiments. But for any given lag-duration the same 8 controls participated in both the object and scene experiments. In addition to the adaptation experiments, all participants completed functional localizer and retinotopy scans, as described below. LSJ and control participants were scanned in multiple sessions at Princeton University. All participants had normal or corrected-to-normal vision and gave informed consent to a protocol approved by Princeton University’s Institutional Review Board.

### fMRI Acquisition and Preprocessing

Participants were scanned in multiple scan sessions with a Siemens Skyra 3T scanner. During each session, high-resolution T1-weighted anatomical scans using the MPRAGE sequence were obtained with the following parameters: TR = 2.3s, TE = 1.97ms, flip angle = 9°, matrix = 256 χ 256, resolution = 1mm isotropic, slices = 176. These anatomical scans were used to align the functional data across sessions. The fMRI scans were acquired with a T2*-weighted echo planar imaging sequence: retinotopy scans, TR = 2.5s, TE = 30ms, flip angle = 75°, matrix = 64 x 64, resolution = 3mm isotropic, slices = 39; functional localizers and adaptation experiments, TR = 2s, TE = 30ms, flip angle = 72°, matrix = 64 x 64, resolution = 3mm isotropic, slices = 36.

The functional scans were preprocessed in AFNI (http://afni.nimh.nih.gov/afni), including de-spiking, slice time correction, motion correction, and de-trending. Although LSJ had significantly more motion during some of the experiments than compared to controls, the amount of motion LSJ had did not differ reliably across different experiments. See Figure S5 for LSJ’s motion parameters for all experiments and for comparison of LSJ and controls’ motion. Thus, any differences between LSJ and controls cannot be merely attributed to differences in motion during scanning.

Data were smoothed with a 4mm full-width half-maximum Gaussian kernel and normalized to percent signal change by dividing the time series by its mean intensity. All functional scans were co-registered to each session’s anatomical scan. FreeSurfer (http://surfer.nmr.mgh.harvard.edu) and SUMA (http://afni.nimh.nih.gov/afni/suma) were used to make inflated and flat cortical surface reconstructions. After preprocessing, the retinotopy and localizer runs were projected onto the inflated brains and voxels that fell within the gray matter boundary were used to define regions-of-interest (ROIs).

### ROI Localization and Retinotopic Mapping

Functional localizers were used to define LOC, PPA, TOS, RSC and FFA in each participant. Runs with alternating blocks of grayscale objects and grid-scrambled objects were used to localize LOC. Separate runs with alternating blocks of grayscale scenes and faces were used to define scene- and face-selective ROIs. Scene-selective ROIs (PPA, TOS, RSC) were defined from scene > face blocks, and the face-selective ROI (FFA) was defined from face > scene blocks. The ROIs were defined using a thresholded *t*-map of *p* < .05, Bonferroni corrected. See Figure 1B for sample stimuli. Each localizer run (2.6 min) began and ended with 8 seconds of fixation. There were eight 12-s blocks in each run, each separated by 6 s of fixation. Each block consisted of 12 different images, randomly selected without replacement from a total of 40 images per category. Stimuli subtended 11° and were presented for 500 ms with an inter-stimulus interval (ISI) of 500 ms. Each participant was scanned in two object localizer runs and two scene/face localizer runs.

Retinotopic areas V1, V2, V3, V4 were defined in each participant across four runs. Each run started with a 10 s fixation period followed by 5 cycles of a wedge that rotated around a central fixation (32 s for a full rotation). The wedge spanned 1-13.5° in eccentricity with an arc length of 45° and was filled with 1000 dots (0.1°, 65 cd/m^2^) moving in random direction at a rate of 7°/s. The wedge rotated either clockwise or counter-clockwise in alternating runs. To delineate visual areas a Fourier analysis was used where the amplitude and phase of the harmonic at the stimulus frequency was determined. The statistical threshold used to delineate ROIs was *p <* 0.001, uncorrected, derived from the *F* ratio of the Fourier transform. Similar phase encoding parameters and procedures were used previously 8, 37, 69) and the details of the statistical analyses were reported previously (70-71).

### Adaptation Experiments

A large set of line drawing objects and colored scene photographs were used for the adaptation experiments. Different sets of object and scene stimuli were used for each experiment and stimuli used for the functional localizers and adaptation experiments did not overlap. Examples are shown in Figure 3A.

#### Immediate adaptation (Experiment 1)

Every participant was scanned in six object and six scene runs. Each run started with 8 s of fixation followed by six blocks each lasting 12 s followed by 12 s of fixation, for a total of 2.5 mins (Figures 3A and 3B). The new and repeat blocks were presented in an alternating fashion. During the new block, 12 distinct objects or scenes were presented. In the repeat block, one object or scene was repeated 12 times. Each image subtended 10° and was presented centrally for 750 ms with an inter-trial-interval of 1s. Square-wave functions matching the timecourse of the design were convolved and regressors for each timepoints for each block were used in a multiple regression model. Additional nuisance regressors included motion parameters, linear drifts within runs, and shifts between runs. The resulting beta values of this regression model were used for the timecourse analyses. To quantify adaptation effects, an adaptation index (AI) was computed for each ROI and participant using similar methods previously published (8, 72):

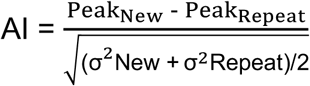

where Peak_New_ and Peak_Repeat_ are defined as the average response over the timepoints at the peak of the hemodynamic response (8-16 s after block onset) for new and repeat blocks, respectively, and σ^2^_New_ and σ^2^_Repeat_ are the variance of these peak responses across new and repeat blocks, respectively. AIs greater than 0 indicate greater responses for new vs. repeat blocks.

#### Long-lag adaptation (Experiments 2-5)

The long-lag adaptation experiments were conducted using a rapid event-related design. Each run started with 8 s of fixation period followed by 36 trials, and ended with 6 s of fixation. During each trial, a stimulus was presented centrally with a fixation cross for 750 ms followed by 250 ms of fixation (Figure 4A). The ITI was 4 s for two-thirds of the trials and 6 s for the remaining one-third of trials, randomized across runs for each subject. During the fixation period of each trial of Experiments 2-4, either a word or a letter prompt (e.g., “A?”) for the alphabet game (described below) was presented for 3 s. There were eight runs for each for the object and scene conditions within every experiment. Repetitions occurred at different lags across experiments: Experiment 2, 30 s (range 2-10 intervening stimuli); Experiment 3, 3 mins (range 36-54 intervening stimuli); Experiment 4 and 5, 6 mins (range 56-70 intervening stimuli). There were 144 new trials and 144 repeat trials for each experiment. Across different experiments, different sets of objects and scenes were used because some of the subjects participated in more than one experiment. BOLD responses were estimated within ROIs using a general linear model with stimulus events and motion parameters from the preprocessing. Stimulus events were modeled with a series of finite impulse response functions, one regressor for each of ten 2-s timepoints. The beta values were converted to percent signal change and averaged over the time period of the peak hemodynamic response (4-8 s after stimulus onset). The difference between the average peak response for the new minus repeat conditions was used to compute AIs for the event-related designs. Since all trials for each condition were used to model the hemodynamic response functions for the new and repeat conditions, these AI computations differed from the AI quantification for the block design experiment, which took into consideration the peak responses for each condition as proportion to the variance across blocks.

### Behavioral Tasks

fMRI runs were designed to be shorter than usual — i.e., no longer than 3 mins, estimated as LSJ’s attention span based on observations and verbal reports from LSJ’s family. LSJ and control participants engaged in a perceptual preference task during the experiments with block designs (i.e., functional localizer and Experiment 1), which encouraged participants to focus on the visual stimuli being presented. After each block of trials, participants were prompted with: “Press a button if you liked what you saw.” To ensure that LSJ did not forget the task during stimulus presentation, task instructions were also shown on the screen during the fixation period before each block: “Now you are going to see some [objects/textures/scenes/faces]. Pay attention!”.

Although this task is subjective in nature, results suggest that LSJ and controls mostly liked the stimuli being presented and responded after most of the blocks. For the localizer runs, controls reported to like the stimuli on 78.1% of the object blocks and 92.2% of the scene blocks. LSJ’s behavioral performance was similar to that of controls (87.5% for both object and scene blocks) and did not differ significantly (*t*_17_ = 0.39, p = and *t*_17_ = −0.29, p = 0.77, respectively). Similarly, for Experiment 1, LSJ (83.3% for objects and 94.4% for scenes) and controls (88.9% for objects and 91.7% for scenes) responded that they liked most of the blocks presented and LSJ’s preference scores did not differ reliably from controls (*t*_7_ = −1.31, p = 0.23 and *t*_7_ = 0.28, p = 0.76, respectively).

While preparing for the study the authors met with LSJ, her family members, and other researchers to assess what kinds of tasks LSJ would be able and motivated to perform, given her limited memory span. On many occasions, we observed that LSJ likes to play an alphabet game where she and another player (e.g., her sister) take turns going through the alphabet and generating a unique word for each letter. With the help of the alphabet game, which provides sequential structure, we observed that LSJ can maintain a fluid conversation for several minutes. Given these observations, we designed a virtual alphabet game between the computer and LSJ that was administered during fMRI. This task encouraged LSJ and control participants to fixate the centrally presented letters and stay engaged during retinotopy and most event-related designs (Experiments 2-4). LSJ and the computer went through the alphabet letter-by-letter, taking turns generating words that begin with the prompted letter. For example, the computer might start by generating a word that starts with an “A” (e.g., “Admire”) and this word is shown at fixation. Then, “B?” would be displayed at fixation and participants needed to generate a word that starts with the letter B.

The words generated on the computer’s turns consisted of mostly non-object words (e.g., verbs and abstract nouns) to avoid interfering with the object and scene stimuli as much as possible and they were not repeated across runs or experiments. Letters and words were displayed on the screen every 4 s. When generating words, participants were instructed to only think of the word and not to say the word out loud, to reduce the possibility of head motion and artifacts related to producing speech. This task allowed us to examine long-lag adaptation when objects/scenes were task-irrelevant.

In Experiment 5, participants performed a go/no-go categorization task on the objects/scenes, to evaluate the role of task relevance. For objects, participants were instructed to press a button if the presented object was a natural object (e.g., plants, animals) and to withhold a response if the object was artificial. For scenes, participants were instructed to press a button if the presented scene was an indoor scene (e.g., kitchen, office) and to withhold a response if the scene was outdoor. Equal numbers of natural and artificial objects and indoor and outdoor scenes were used.

### Single Case Study Statistics

The statistical comparison of LSJ to the control group was done with a modified independent samples two-tailed *t*-test (38), which accounts for the limited size of the control group and tests the null hypothesis that the single case comes from a population of controls. This method has been widely used in neuropsychological case studies (8, 73-74) and has advantages over other single case statistics such as the modified ANOVA or z-score inferences (75). As a visual benchmark, we compared LSJ’s results to a representative control participant C1 for each experiment. We confirmed that C1 was representative by comparing their data (e.g., size of ROIs, amplitude of adaptation index) to the rest of the control group for each experiment. None of the tests resulted in a significant difference (all *p*s > 0.05).

## Acknowledgements

We thank LSJ’s sister for her extensive help and feedback, and dedicate this article to the memory of their mother, who also provided considerable support. This research was funded by NIH EY02316601 (JGK); Johns Hopkins Brain Science Institute (BL & MM); NIH R01 EY021755 (NTB); NSF BCS 1025149 (SK); NIH 2RO1 MH64043 (SK).

## Supplemental Information

**Figure S1.**
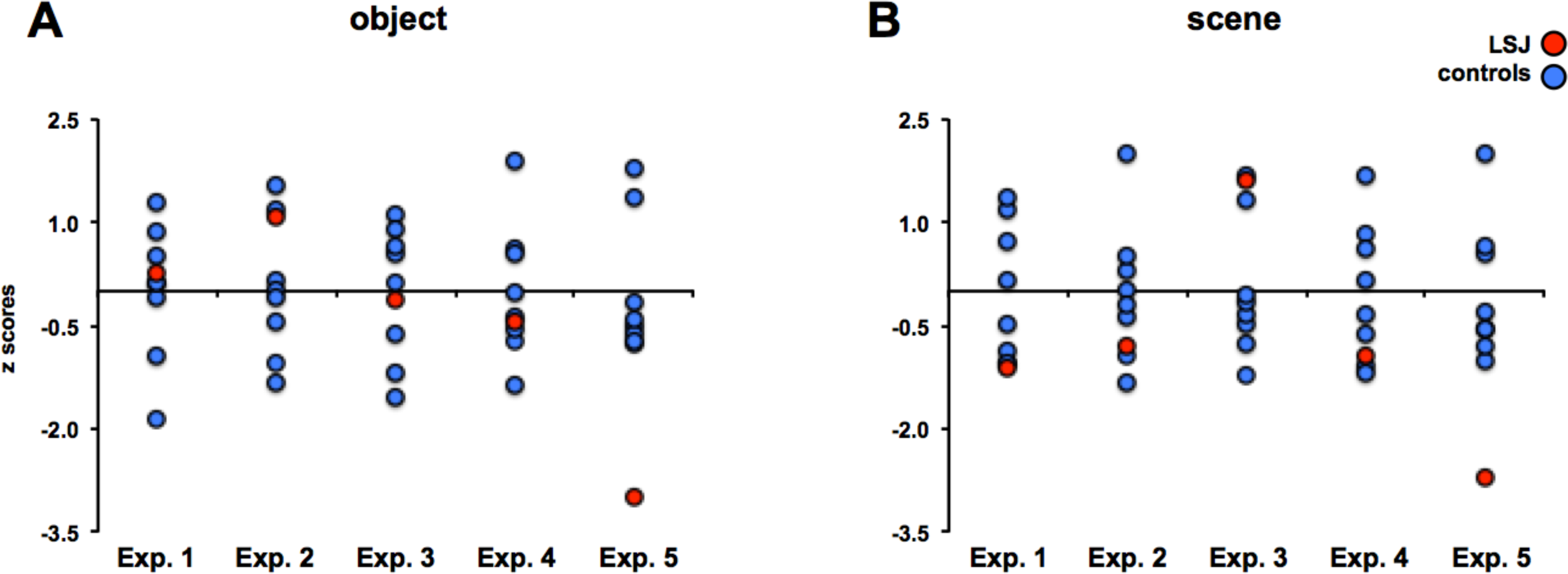
AI comparisons between LSJ and individual controls. Normalized (z-scores with mean and SD from controls) AIs from Experiments 1-5 are plotted for LSJ (in red) and controls (in blue) for objects (A) and scenes (B). For Exps. 1-4, LSJ’s AIs are within the distribution of controls’ AIs. In Exp. 5, for both object and scene, LSJ’s AI values were about 3 SD below control mean AI values.

**Figure S2.**
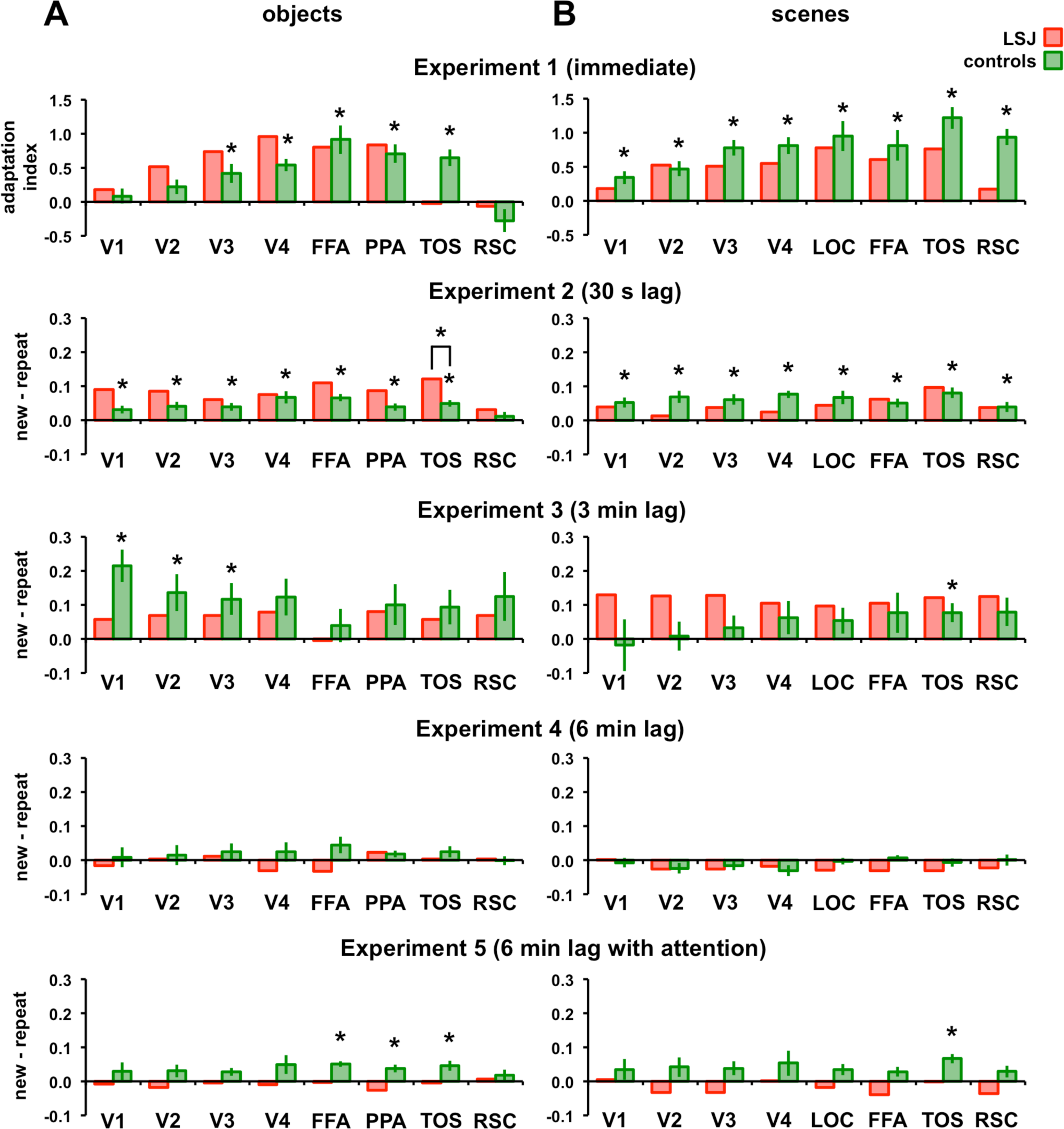
Adaptation effects in visual ROIs across experiments. AIs from Experiments 1-5 are plotted for LSJ (in red) and controls (in green) for visual ROIs for objects (A) and scenes (B). For this and subsequent figures and tables, one-sample t-tests were used to determine reliably positive AIs for controls (* p < 0.05) and Crawford and Howell’s t-test (1) for a single case (two-tailed) was used to compare AIs between LSJ and controls. The t-values and p-values for reliable adaptation effects in these ROIs for controls and the comparison between LSJ and controls are reported in Table S1 for objects and Table S2 for scenes.

**Table S1.** T-values and p-values (in parentheses) for adaptation effects in visual ROIs across experiments for objects in controls, and the comparison bewteen LSJ and controls.

|  |  | V1 | V2 | V3 | V4 | LOC | FFA | TOS | RSC |
| --- | --- | --- | --- | --- | --- | --- | --- | --- | --- |
| **Exp. 1**<br>(immediate) | **Controls** | 0.75 (0.48) | 2.00 (0.08) | 2.98 (0.02) | 5.97 (< 0.01) | 4.34 (< 0.01) | 5.17 (< 0.01) | 5.26 (< 0.01) | -1.67 (0.14) |
|  | **LSJ vs. Controls** | 0.30 (0.77) | 0.88 (0.41) | 0.77 (0.47) | 1.55 (0.16) | -0.17 (0.87) | 0.32 (0.76) | -1.81 (0.11) | 0.43 (0.68) |
| **Exp. 2**<br>(30 s) | **Controls** | 2.75 (0.03) | 3.29 (0.01) | 3.34 (0.01) | 3.65 (< 0.01) | 5.91 (< 0.01) | 3.71 (< 0.01) | 5.08 (< 0.01) | 0.91 (0.39) |
|  | **LSJ vs. Controls** | 1.76 (0.12) | 1.17 (0.28) | 0.58 (0.58) | 0.14 (0.89) | 1.28 (0.24) | 1.56 (0.16) | 2.48 (0.04) | 0.49 (0.64) |
| **Exp. 3**<br>(3 min) | **Controls** | 4.49 (< 0.01) | 2.51 (0.04) | 2.52 (0.04) | 2.28 (0.06) | 0.79 (0.46) | 1.68 (0.14) | 1.83 (0.11) | 1.74 (0.12) |
|  | **LSJ vs. Controls** | -1.09 (0.31) | -0.41 (0.69) | -0.35 (0.74) | -0.27 (0.79) | -0.32 (0.76) | -0.11 (0.92) | -0.23 (0.82) | -0.26 (0.80) |
| **Exp. 4**<br>(6 min) | **Controls** | 0.28 (0.78) | 0.50 (0.63) | 0.95 (0.37) | 0.82 (0.44) | 1.82 (0.11) | 1.74 (0.13) | 1.49 (0.18) | -0.16 (0.88) |
|  | **LSJ vs. Controls** | -0.28 (0.79) | -0.14 (0.89) | -0.17 (0.87) | -0.64 (0.54) | -1.04 (0.33) | 0.15 (0.88) | -0.45 (0.67) | 0.15 (0.88) |
| **Exp. 5**<br>(6 min attention) | **Controls** | 1.14 (0.29) | 1.63 (0.15) | 2.30 (0.06) | 1.78 (0.12) | 5.84 (< 0.01) | 3.40 (0.01) | 3.05 (0.02) | 1.01 (0.35) |
|  | **LSJ vs. Controls** | -0.48 (0.65) | -0.87 (0.41) | -0.92 (0.39) | -0.71 (0.50) | -2.08 (0.08) | -1.92 (0.10) | -1.13 (0.29) | -0.20 (0.85) |

**Table S2.** T-values and p-values (in parentheses) for adaptation effects in visual ROIs across experiments for scenes in controls, and the comparison between LSJ and controls.

|  |  | V1 | V2 | V3 | V4 | LOC | FFA | TOS | RSC |
| --- | --- | --- | --- | --- | --- | --- | --- | --- | --- |
| **Exp. 1**<br>(immediate) | **Controls** | 3.53 (0.01) | 4.31 (< 0.01) | 6.72 (< 0.01) | 6.55 (< 0.01) | 4.32 (< 0.01) | 3.58 (0.01) | 7.66 (< 0.01) | 7.73 (< 0.01) |
|  | **LSJ vs. Controls** | -0.55 (0.60) | 0.16 (0.87) | -0.79 (0.46) | -0.70 (0.50) | -0.26 (-0.80) | -0.30 (0.77) | -0.95 (0.38) | -2.10 (0.07) |
| **Exp. 2**<br>(30 s) | **Controls** | 3.48 (0.01) | 3.93 (0.01) | 3.89 (0.01) | 7.00 (< 0.01) | 3.45 (0.01) | 4.00 (0.01) | 5.15 (< 0.01) | 2.63 (0.03) |
|  | **LSJ vs. Controls** | -0.27 (0.79) | -1.06 (0.32) | -0.49 (0.64) | -1.56 (0.16) | -0.40 (0.70) | 0.33 (0.76) | 0.34 (0.75) | 0.37 (0.72) |
| **Exp. 3**<br>(3 min) | **Controls** | -0.24 (0.82) | 0.18 (0.86) | 0.94 (0.38) | 1.28 (0.24) | 1.40 (0.20) | 1.29 (0.24) | 2.78 (0.03) | 1.91 (0.10) |
|  | **LSJ vs. Controls** | 0.65 (0.54) | 1.31 (0.23) | 0.92 (0.39) | 0.28 (0.79) | 0.37 (0.72) | 0.15 (0.88) | 0.54 (0.60) | 0.37 (0.73) |
| **Exp. 4**<br>(6 min) | **Controls** | -0.58 (0.58) | -1.55 (0.17) | -1.24 (0.25) | -1.85 (0.11) | -0.33 (0.75) | 0.80 (0.45) | -0.62 (0.55) | 0.01 (0.99) |
|  | **LSJ vs. Controls** | 0.22 (0.84) | -0.03 (0.97) | -0.23 (0.82) | 0.24 (0.82) | -0.89 (0.40) | -1.50 (0.18) | -0.66 (0.53) | -0.47 (0.65) |
| **Exp. 5**<br>(6 min attention) | **Controls** | 1.14 (0.29) | 1.45 (0.19) | 1.83 (0.11) | 1.42 (0.20) | 2.12 (0.07) | 1.96 (0.09) | 4.85 (< 0.01) | 1.64 (0.15) |
|  | **LSJ vs. Controls** | -0.33 (0.75) | -0.87 (0.41) | -1.13 (0.30) | -0.46 (0.66) | -1.07 (0.32) | 0.27 (0.80) | -1.62 (0.15) | -1.22 (0.26) |

**Figure S3.**
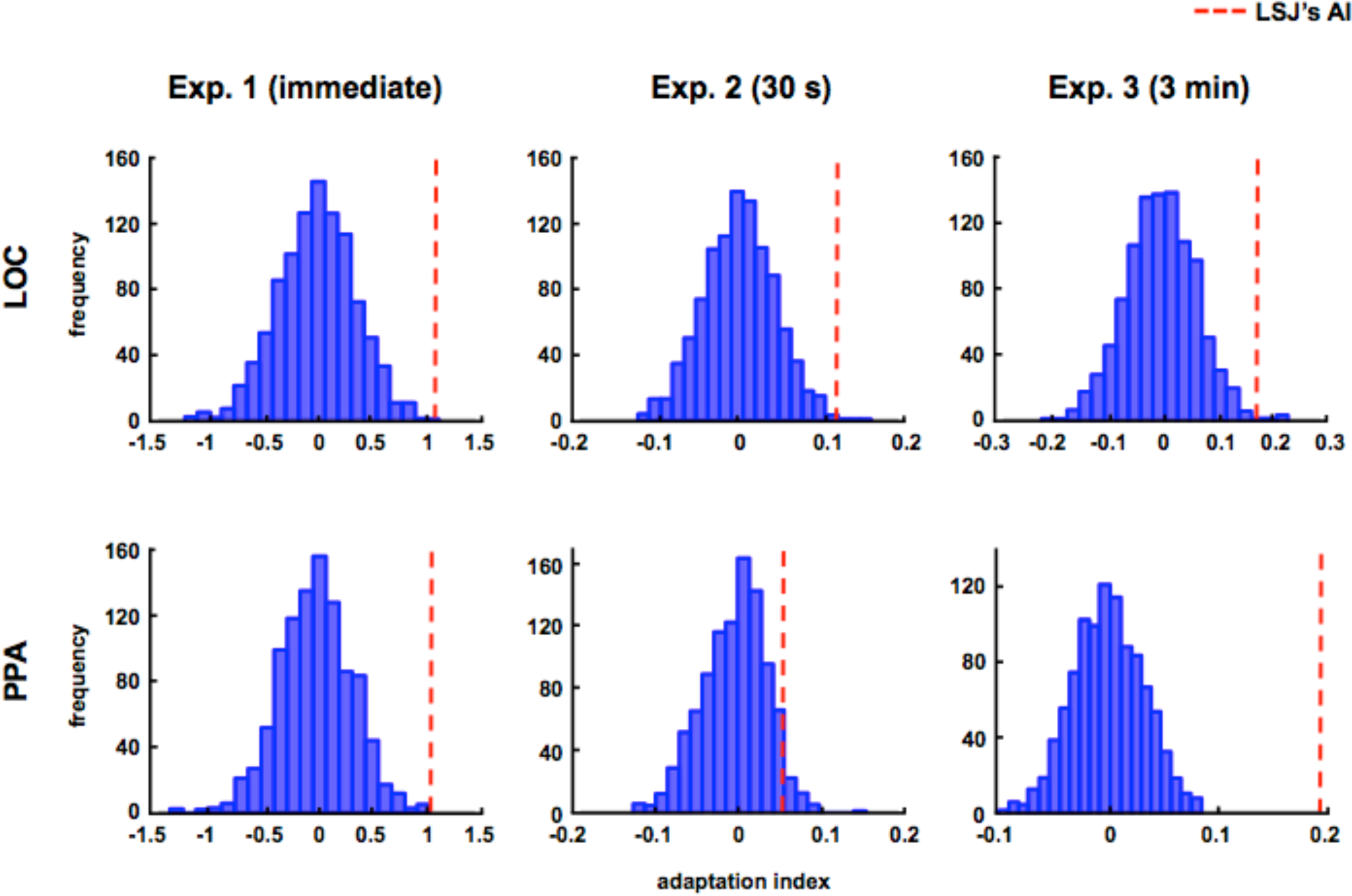
Permutation test for LSJ’s adaptation effects. To examine whether or not LSJ’s adaptation effects are expected by chance, we performed a permutation test to compute the null distribution of adaptation indices (AI). We randomly assigned condition labels to each block (Exp. 1) or trial (Exp. 2 and Exp. 3) using LSJ’s data and computed AIs over 1000 iterations. The distributions of the null AIs are plotted in blue for each experiment and for each ROI where LSJ showed adaptation. The red dashed vertical line indicates LSJ’s AIs as a reference. In all of the experiments, LSJ’s AI was within the 5% tail of the distribution (Exp. 1, LOC: 0.1%, PPA: 0.1%; Exp. 2, LOC: 0.3%, PPA: 4.5%; Exp. 3, LOC: 0.9%, PPA: 0%).

**Figure S4.**
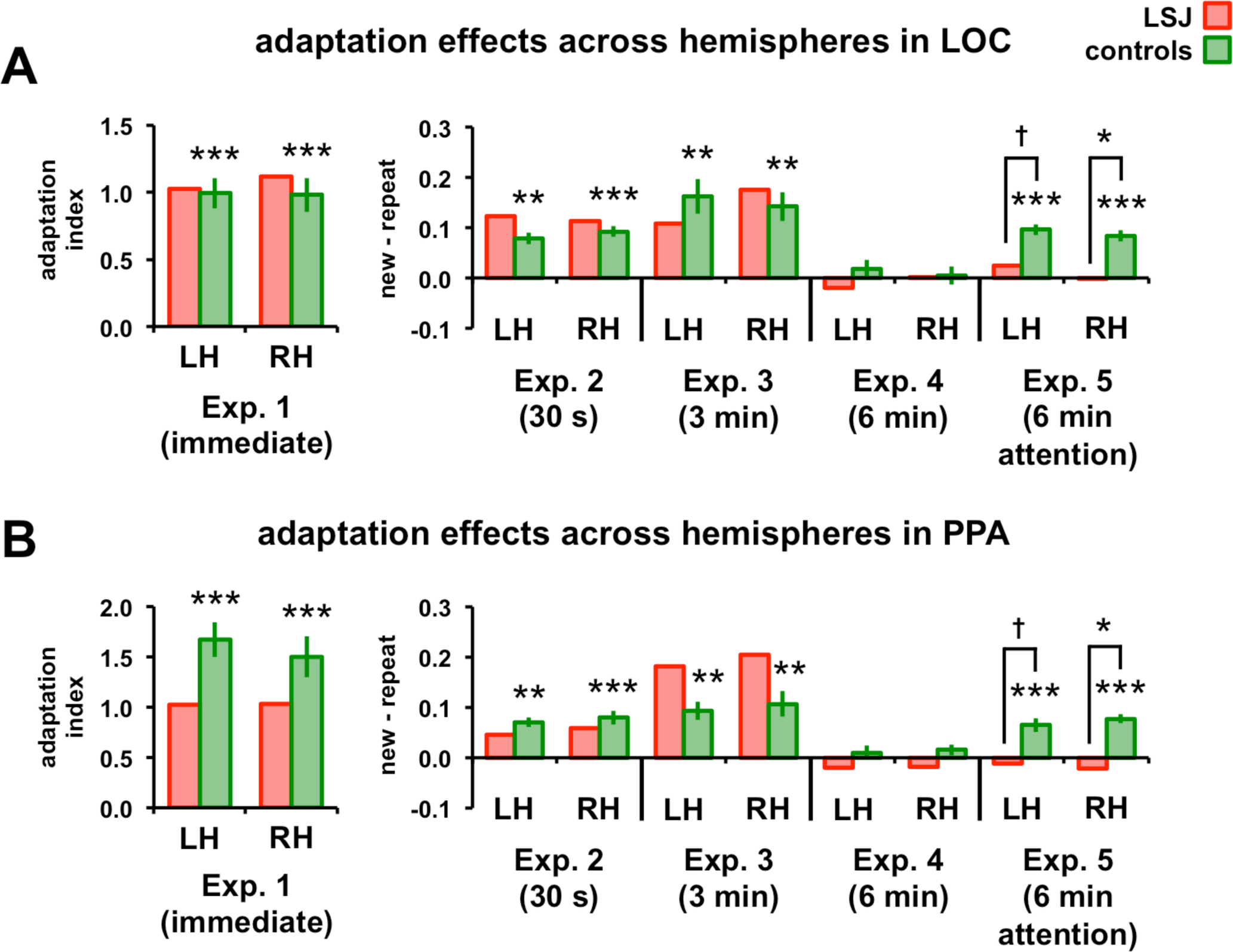
Adaptation effects by hemisphere across experiments. AIs for left and right LOC for objects (A) and PPA for scenes (B) are plotted across experiments. Controls showed reliably positive AIs in both hemispheres for all experiments except for Experiment 4 (*** p < 0.001, ** p < 0.01). In Experiment 5, LSJ showed marginally reduced AIs in left († p ≤ 0.1) and significantly in the right (* p < 0.05) hemispheres for both LOC and PPA. The t-values and p-values for reliable adaptation effects in left and right hemispheres in controls and the comparison between LSJ and controls are reported in Table S3 for objects and Table S4 for scenes.

**Table S3.** T-values and p-values (in parentheses) for adaptation effects in left and right LOC across experiments for objects in controls, and the comparison between LSJ and controls.

|  |  | LOC LH | LOC RH | LH vs RH |
| --- | --- | --- | --- | --- |
| **Exp. 1**<br>(immediate) | **Controls** | 8.75 (< .01) | 7.81 (< 0.01) | 0.21 (0.84) |
|  | **LSJ vs. Controls** | 0.09 (0.93) | 0.36 (0.73) | -0.53 (0.61) |
| **Exp. 2**<br>(30 s) | **Controls** | 6.64 (< .01) | 8.42 (< 0.01) | -2.33 (0.05) |
|  | **LSJ vs. Controls** | 1.27 (0.25) | 0.64 (0.54) | 1.34 (0.22) |
| **Exp. 3**<br>(3 min) | **Controls** | 4.61 (< .01) | 4.92 (< 0.01) | 0.91 (0.39) |
|  | **LSJ vs. Controls** | -0.52 (0.62) | 0.39 (0.71) | -1.33 (0.22) |
| **Exp. 4**<br>(6 min) | **Controls** | 1.03 (0.34) | 0.25 (0.81) | 1.05 (0.33) |
|  | **LSJ vs. Controls** | -0.73 (0.49) | -0.07 (0.95) | -0.92 (0.39) |
| **Exp. 5**<br>(6 min attention) | **Controls** | 8.98 (< 0.01) | 7.40 (< 0.01) | 2.35 (0.05) |
|  | **LSJ vs. Controls** | -2.21 (0.06) | -2.51 (0.04) | 0.98 (0.36) |

**Table S4.**
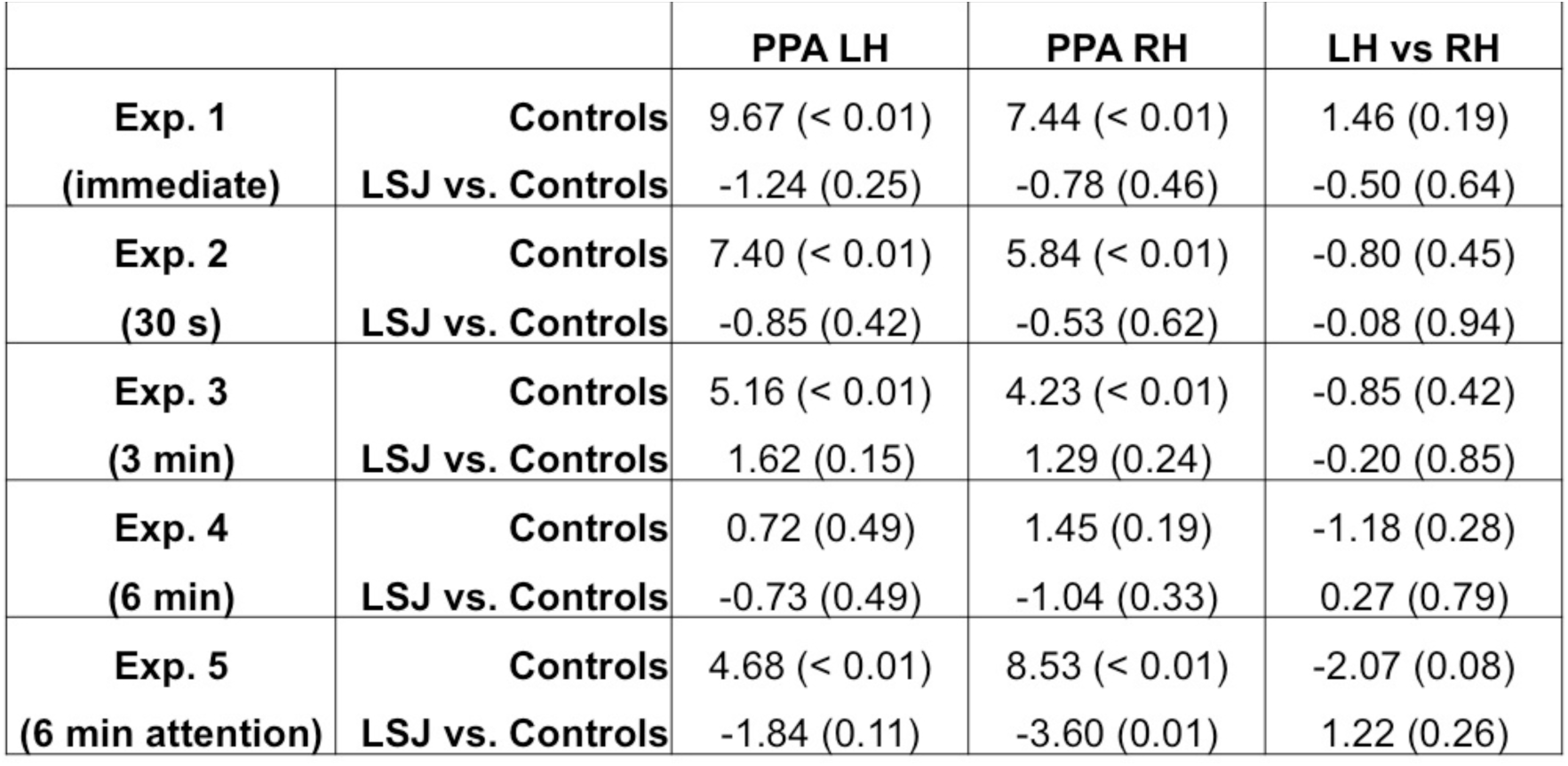
T-values and p-values (in parenthesis) for adaptation effects in left and right PPA across experiments for scenes in controls, and the comparison between LSJ and controls.

**Figure S5.**
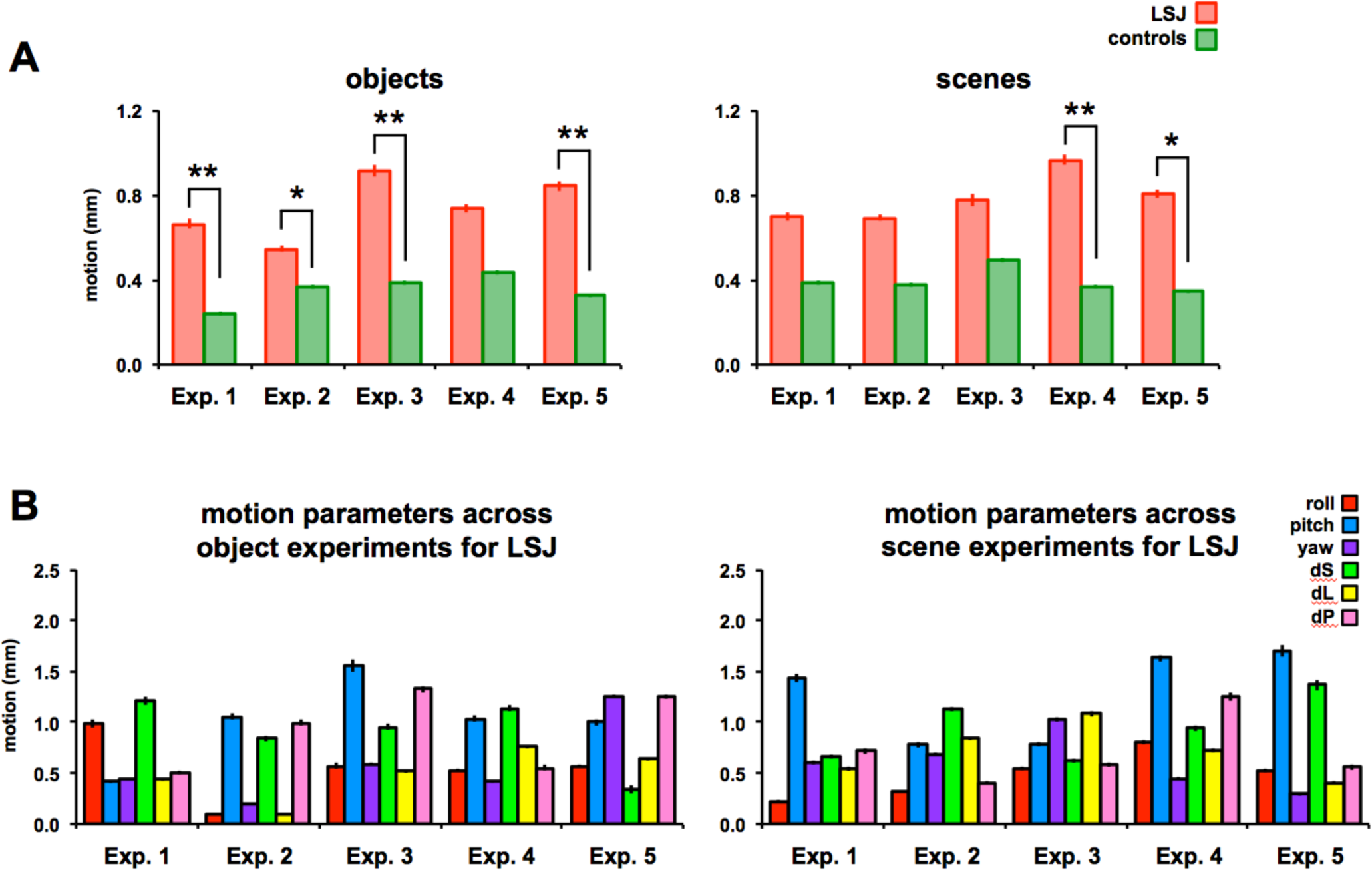
Comparison of motion parameters between LSJ and controls and across experiments for LSJ. (A) The six motion directions (roll, pitch, yaw, dS, dL and dP) were averaged for each subject for each experiment. LSJ had significantly more motion than controls in 6/10 experiments (* p < 0.05, ** p < 0.01). However, the amount of motion for LSJ did not differ across experiments (B) as assessed by a two-way ANOVA with experiments (1-5) and motion direction (roll, pitch, yaw, dS, dL, and dP) as independent variables. There was no main effect of experiment for both objects (*F*_4,20_ = 1.02, p = 0.42) and scenes (*F*_4,20_ = 0.69, p = 0.61).

**Figure S6.**
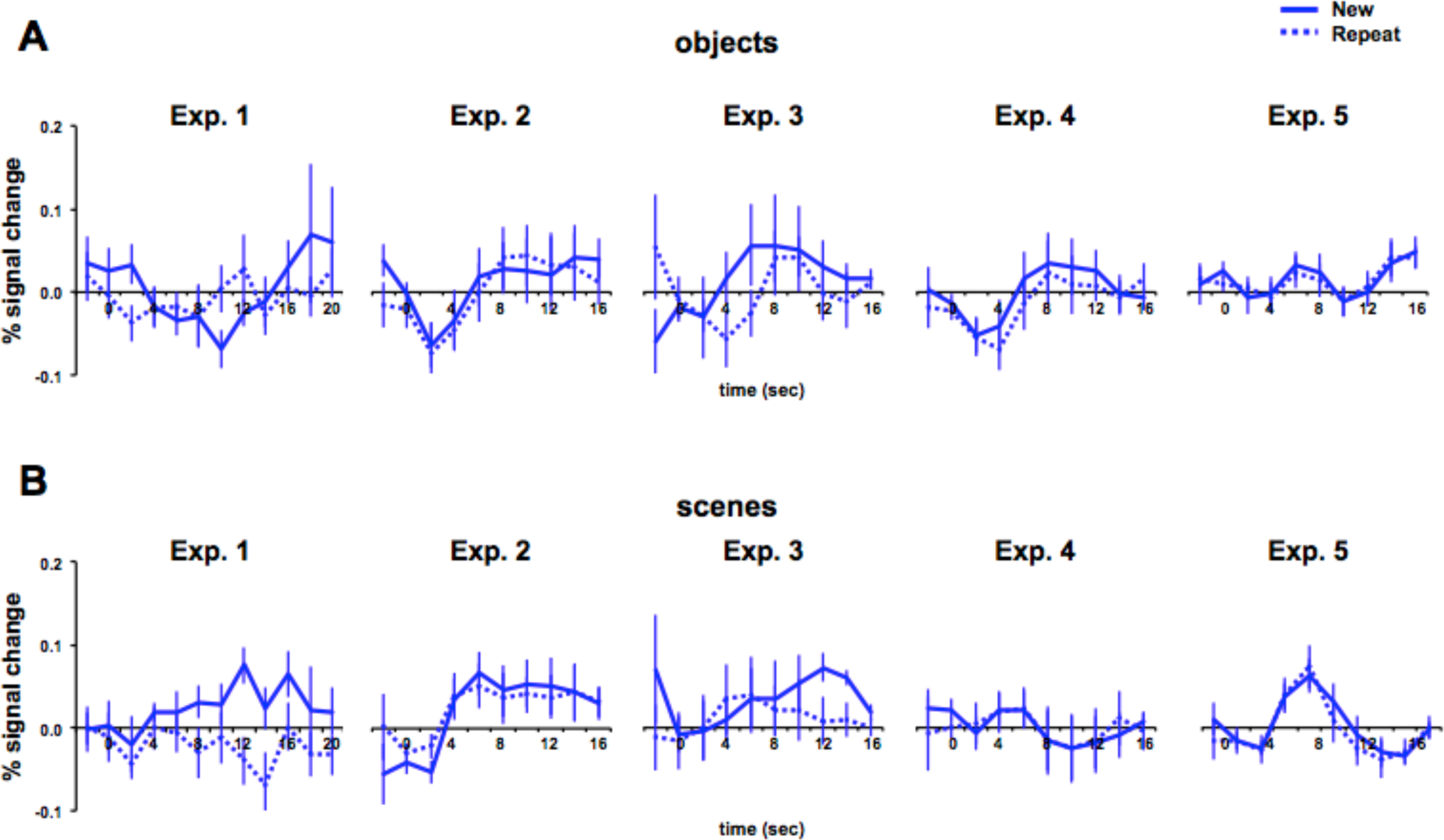
Hippocampus activity across experiments for controls. Hippocampus was anatomically localized using Freesurfer’s (http://surfer.nmr.mgh.harvard.edu) subcortical segmentation tool (2) for each control subject. Hippocampus activity was examined across all object (A) and scene (B) experiments. During Exp. 1, there was no visually evoked responses in the hippocampus for both object and scene tasks. For comparison, refer to Figure 3 for visually evoked responses in LOC and PPA for Exp. 1. In Exp. 5, there are clear evoked responses in the hippocampus (especially for scenes) suggesting that the hippocampus is involved in the long-lag adaptation with directed attention which supports the impairment in LSJ.

## References

1. Grill-Spector K, Kushnir T, Edelman S, Avidan G, Itzchak Y, Malach R (1999) Differential processing of objects under various viewing conditions in the human lateral occipital complex. Neuron 24:187–203.

2. Grill-Spector K, Malach R (2001) FMR-adaptation: A tool for studying the functional properties of human cortical neurons. Acta Psychol 107:293–321.

3. Hatfield M, McCloskey M, Park S (2016) Neural representation of object orientation: A dissociation between MVPA and repetition suppression. Neuroimage 139:136–148.

4. Kim JG, Biederman I, Lescroart MD, Hayworth KJ (2009) Adaptation to objects in the lateral occipital complex (LOC): Shape or semantics? Vision Res 49:2297–2305.

5. Konen CS, Kastner S (2008) Two hierarchically organized neural systems for object information in human visual cortex. Nat Neurosci 11:224–231.

6. Epstein R, Harris A, Stanley D, Kanwisher N (1999) The parahippocampal place area: recognition, navigation, or encoding? Neuron 23:115–125.

7. Epstein RA, Parker WE, Feiler AM (2008) Two kinds of fMRI repetition suppression? Evidence for dissociable neural mechanisms. J Neurophys 99:2877–2886.

8. Kim JG, Aminoff EM, Kastner S, Behrmann M (2015) A neural basis for developmental topographic disorientation. J Neurosci 35:12954–12969.

9. van Turennout M, Ellmore T, Martin A (2000) Long-lasting cortical plasticity in the object naming system. Nature 3:1329–1334.

10. van Turennout M, Bielamowicz L, Martin A (2003) Modulation of neural activity during object namging: Effects of time and practice. Cereb Cortex 13:381–391.

11. Henson R, Shallice T, Dolan R (2000) Neuroimaging evidence for dissociable forms of repetition priming. Science 287:1269–1272.

12. Henson RN, Ross ARE, Vuilleumeir P, Rugg MD (2004) The effect of repetition lag on electrophysiological and haemodynamic correlates of visual object priming. NeuroImage 21:1674–1689.

13. Weiner KS, Sayres R, Winberg J, Grill-Spector K (2010) FMRI-adaptation and category selectivity in human ventral temporal cortex: Regional differences across time scales. J Neurophys 103:3349–3365.

14. Brozinsky CJ, Yonelinas AP, Kroll NEA, Ranganath C (2005) Lag-sensitive repetition suppression effects in the anterior parahippocampal gyrus. Hippocampus 15:557–561.

15. Grill-Spector K, Henson R, Martin A (2006) Repetition and the brain: Neural models of stimulus-specific effects. Trends Cog Sci 10:14–23.

16. Norman KA, O’Reilly RC (2003) Modeling hippocampal and neocortical contributions to recognition memory: A contemporary-learning-systems approach. Psych Rev 110:611–646.

17. Felleman DJ, Van Essen DC (1991) Distributed hierarchical processing in the primate cerebral cortex. Cereb Cortex 1:1–47.

18. Turk-Browne NB, Scholl BJ, Johnson MK, Chun MM (2010) Implicit perceptual anticipation triggered by statistical learning. J Neurosci 30:11177–11187.

19. Staresina BP, Henson RNA, Kriegeskorte N, Alink A (2012) Episodic reinstatement in the medial temporal lobe. J Neurosci 32:18150–18156.

20. Bosch SE, Jehee JFM, Fernandez G, Doeller CF (2014) Reinstatement of associative memories in early visual cortex is signaled by the hippocampus. J Neurosci 34:7493–7500.

21. O’Reilly RC, Rudy JW (2001) Conjunctive representation in learning and memory: Principles of cortical and hippocampal function. Psych Rev 108:311–345.

22. Rolls ET, Cahusac PMB, Feigenbaum JD, Miyashita Y (1993) Responses of single neurons in the hippocampus of the macaque related to recognition memory. Exp Brain Res 93:299–306.

23. Vannini P, Hedden T, Sullivan C, Sperling RA (2013) Differential functional response in the posteromedial cortices and hippocampus to stimulus repetition during successful encoding. Hum Brain Mapp 34:1568–1578.

24. Kremers NAW, Deuker L, Kranz TA, Oehrn C, Fell J, Axmacher N (2014) Hippocampal control of repetition effects for associative stimuli. Hippocampus 24:892–902.

25. Gregory E, McCloskey M, Landau B (2014) Profound loss of general knowledge in retrograde amnesia: evidence from an amnesic artist. Front Hum Neurosci 8:287.

26. Gregory E, McCloskey M, Ovans Z, Landau B (2016) Declarative memory and skill-related knowledge: Evidence from a case of amnesia and implications for theories of memory. Cog Neuropsychol 33:220–240.

27. Schapiro AC, Gregory E, Landau B, McCloskey M, Turk-Browne NB (2014) The necessity of the medial temporal lobe for statistical learning. J Cogn Neurosci 26:1736–1747.

28. Valtonen J, Gregory E, Landau B, McCloskey M (2014) New learning of music after bilateral medial temporal lobe damage: Evidence from an amnesic patient. Front Hum Neurosci 8:694.

29. Aly M, Turk-Browne NB (2016b) Attention stabilizes representations in the human hippocampus. Cereb Cortex 26:783–796.

30. Uncapher MR, Rugg MD (2009) Selecting for memory? The influence of selective attention on the mnemonic binding of contextual information. J Neurosci 29:8270–8279.

31. Carr VA, Engel SA, Knowlton BJ (2013) Top-down modulation of hippocampal encoding activity as measured by high-resolution functional MRI. Neuropsychologia 51:1829–1837.

32. Aly M, Turk-Browne NB (2016a) Attention promites episodic encoding by stabilizing hippocampal representations. Proc Natl Acad Sci USA 113:E420–429.

33. Murray SO, Wojciulik E (2003) Attention increases neural selectivity in the human lateral occipital complex. Nat Neurosci 7:70–74.

34. Moore KS, Yi D-J, Chun M (2013) The effect of attention on repetition suppression and multivoxel pattern similarity. J Cog Neurosci 25:1305–1314.

35. Eger E, Henson RNA, Driver J, Dolan RJ (2004) BOLD repetition decreases in object-responsive ventral visual areas depend on spatial attention. J Neurophysiol 92:1241–1247.

36. Yi D-J, Chun MM (2005) Attentional modulation of learning-related repetition attenuation effects in human parahippocampal cortex. J Neurosci 25:3593–3600.

37. Bandettini PA, Jesmanowicz A, Wong EC, Hyde JS (1993) Processing strategies for time-course data sets in functional MRI of the human brain. Magn Reson Med 30:161–173.

38. Crawford JR, Howell DC (1998) Regression equations in clinical neuropsychology: An evaluation of statistical methods for comparing predicted and obtained scores. J Clin Exp Neuropsychol 20:755–762.

39. Meister IG, Weidemann J, Foltys H, Brand H, Willmes K, Krings T, Thron A, Topper R, Boroojerdi B (2005) The neural correlate of very-long-term picture priming. Eu. J Neurosci 21:1101–1106.

40. Koutstaal W, Wagner AD, Rotte M, Maril A, Buckner RL, Schacter DL (2001) Perceptual specificity in visual object priming: Functional magnetic resonance imaging evidence for a laterality difference in fusiform cortex. Neuropsychologia 39:184–199.

41. Dobbins IG, Schnyer DM, Verfaellie M, Schacter D (2004) Cortical activity reductions during repetition priming can restult from rapid response learning. Nature 428:316–319.

42. Turk-Browne NB, Yi D-J, Chun MM (2006) Linking implicit and explicit memory: Common encoding factors and shared representations. Neuron 49:917–927.

43. Vuilleumier P, Schwartz S, Duhoux S, Dolan RJ, Driver J (2006) Selective attention modulates neural substrates of repetition priming and “implicit” visual memory: Suppressions and enhancements revealed by fMRI. J Cogn Neurosci 17:1245–1260.

44. Warrington EK, Weiskrantz L (1974) The effect of prior learning on subsequent retention in amnesic patients. Neuropsychologia 12:419–428.

45. Gabrieli JDE, Milberg W, Keane MM, Corkin S (1990) Intact priming of patterns despite impaired memory. Neuropsychologia 28:417–427.

46. Cave BC, Squire LR (1992) Intact and long-lasting repetition priming in amnesia. J Exp Psychol 18:509–520.

47. Maccotta L, Buckner RL (2004) Evidence for neural effects of repetition that directly correlate with behavioral priming. J Cog Neurosci 16:1625–1632.

48. McMahon DBT, Olson CR (2007) Repetition suppression in monkey inferotemporal cortex: Relation to behavioral priming. J Neurophysiol 97:3532–3543.

49. Nagy ME, Rugg MD (1989) Modulation of event-related potentials by word repetition: The effects of inter-item lag. Psychophysiol 26:431–436.

50. Li L, Miller EK, Desimone R (1993) The representation of stimulus familiarity in anterior inferior temporal cortex. J Neurophys 69:1918–1929.

51. Anderson B, Mruczek, REB, Kawasaki K, Sheinberg D (2008). Effects of familiarity on neurla activity in monkey inferior temporal lobe. Cereb Cortex 18:2540–2552.

52. Freedman DJ, Riesenhuber M, Poggio T, Miller EK (2006) Experience-dependent sharpening of visual shape selectivity in inferior temporal cortex. Cereb Cortex 16:1631–1644.

53. Hasson U, Chen J, Honey CJ (2015) Hierarchical process memory: Memory as an integral compoent of information processing. Trends Cogn Sci 19:304–313.

54. Xiang J-X, Brown MW (1998) Differential neuronal encoding of novelty, familiarity and recency in regions of the anterior temporal lobe. Neuropharmacol 37:657–676.

55. Fahy FL, Riches IP, Brown MW (1993) Neuronal activity related to visual recognition memory: Long-term memory and the encoding of recency and familiarity information in the primate anterior and medial inferior temporal and rhinal cortex. Exp Brain Res 96:457–472.

56. Gordon AM, Rissman J, Kiani R, Wagner AD (2014) Cortical reinstatement mediates the relationship between content-specific encoding activity and subsequent recollection decisions. Cereb Cortex 24:3350–3364.

57. Danker JF, Tompary A, Davachi L (2017) Trial-by-trial hippocampal encoding activation predicts the fidelity of cortical reinstatement during subsequent retrieval. Cereb Cortex 27:3515–3524.

58. Manelis A, Wheeler ME, Paynter CA, Storey L, Reder LM (2011) Opposing patterns of neural priming in same-exemplar vs. different-exemplar repetition predict subsequent memory. NeuroImage 55:763–772.

59. Smith CN, Squire LR (2007) Experience-dependent eye movements reflect hippocampus-dependent (aware) memory. J Neurosci 28:12825–12833.

60. Smith CN, Squire LR (2017) When eye movements express memory for old and new scenes in the absence of awareness and independent of hippocampus. Learn & Mem 24:95–103.

61. Chun MM, Turk-Browne NB (2007) Interactions between attention and memory. Curr Opin Neurobiol 17:177–184.

62. Summerfield C, de Lange FP (2014) Expectation in perceptual decision making: Neural and computational mechanisms. Nat Rev Neurosci 15:745–756.

63. Bar M, Kassam KS, Ghuman AS, Boshyan J, Schmid AM, Dale AM, Hamalainen, MS, Marinkovic K, Schacter DL, Rosen BR, Halgren E (2006) Top-down facilitation of visual recognition. Proc Natl Acad Sci USA 103:449–454.

64. O’Doherty J, Dayan P, Schultz J, Deichmann R, Friston K, Dolan RJ (2004) Dissociable roles of ventral and dorsal striatum in instrumental conditioning. Science 304:452–454.

65. Schapiro AC, Turk-Browne NB, Botvinick MM, Norman KA (2017) Complementary learning systems within the hippocampus: A neural network modelling approach to reconciling episodic memory with statistical learning. Phil Trans R Soc Lond B Biol Sci 372:1711.

66. Hindy NC, Ng FY, Turk-Browne NB (2016) Linking pattern completion in the hippocampus to predictive coding in visual cortex. Nat Neurosci 19:665–667.

67. Wig GS, Grafton ST, Demos KE, Kelley WM (2005) Reductions in neural activity underlie behavioral components of repetition priming. Nat Neurosci 8:1228–1135.

68. Xu Y, Turk-Browne NB, Chun MM (2007) Dissociating task performance from fMRI repetition attenuation in ventral visual cortex. J Neurosci 27:5981–5985.

69. Swisher JD, Halko MA, Merabet LB, McMains SA, Somers DC (2007) Visual topography of human intraparietal sulcus. J Neurosci 27:5326–5337.

70. Schneider KA, Richter MC, Kastner S (2004) Retinotopic organization and functional subdivisions of the human lateral geniculate nucleus: A high-resolution functional magnetic resonance imaging study. J Neurosci 24:8975–8985.

71. Arcaro MJ, McMains SA, Singer BD, Kastner S (2009) Retinotopic organization of human ventral visual cortex. J Neurosci 29:10638–10652.

72. Pinsk MA, Arcaro M, Weiner KS, Kalkus JF, Inati SJ, Gross CG, Kastner S (2009) Neural representations of faces and body parts in macaque and human cortex: A comparative fMRI study. J Neurophysiol 101:2581–2600.

73. Behrmann M, Peterson MA, Moscovitch M, Suzuki S (2006) Independent representation of parts and the relations between them: Evidence from integrative agnoisa. J Exp Psychol 32:1169–1184.

74. Konen CS, Behrmann M, Nishimura M, Kastner S (2011) The functional neuroanatomy of object agnosia: A case study. Neuron 71:49–60.

75. Crawford JR, Garthwaite PH, Howell DC, Gray CD (2004) A single case with a control sample: Modified t-tests versus Mycroft et al.’s (2002) modified ANOVA. Cog Neuropsych 21:750–755.

## References

1. Crawford JR, Howell DC (1998) Regression equations in clinical neuropsychology: An evaluation of statistical methods for comparing predicted and obtained scores. J Clin Exp Neuropsychol 20:755–762.

2. Fischl B, Salat DH, Busa E, Albert M, Dieterich M, Haselgrove C, van der Kouwe A, Killiany R, Kennedy D, Klaveness S, Montillo A, Makris N, Rosen B, Dale AM (2002) Whole brain segmentation: Automated labeling of neuroanatomical structures in the human brain. Neuron 33:341–355.

